# TiMEG: an integrative approach for partially missing multi-omics data with an application to tuberous sclerosis

**DOI:** 10.1101/2020.12.10.420638

**Authors:** Sarmistha Das, Indranil Mukhopadhyay

## Abstract

Multi-omics data integration is widely used to understand the genetic architecture of disease. In multi-omics association analysis, data collected on multiple omics for the same set of individuals are immensely important for biomarker identification. But when the sample size of such data is limited, the presence of partially missing individual-level observations poses a major challenge in data integration. More often, genotype data are available for all individuals under study but gene expression and/or methylation information are missing for different subsets of those individuals. Here, we develop a statistical model TiMEG, for the identification of disease-associated biomarkers in a case-control paradigm by integrating the above-mentioned data types, especially, in presence of missing omics data. Based on a likelihood approach, TiMEG exploits the inter-relationship among multiple omics data to capture weaker signals, that remain unidentified in single-omics analyses. Its application on a real tuberous sclerosis dataset identified functionally relevant genes in the disease pathway.

## 2 Introduction

Advances in next-generation sequencing (NGS) technologies have led to harness robust structural and functional knowledge of the human genome. Thus, NGS provides unprecedented opportunities to understand health and disease at the present time^1–4^. But the transformation of knowledge from bench to bedside lies in mining the massively available data for genes and variants of high clinical relevance^5^. To understand the genetic architecture of disease, genome-wide association studies^6,7^ and several other studies based on single-omics data such as gene expression or DNA methylation have cataloged many disease-associated loci. Nonetheless, we are yet to understand the etiology of many complex diseases as they occur due to an intricate interplay of various genetic elements^8^.

Multi-omics data integration serves as a springboard for unravelling such inherent complexity underlying the genetic architecture of a disease^8,9^. Data integration through statistical and/or computational models, plays a major role in the prediction of genomic and environmental perturbations underlying disease/complex traits, and transferring preclinical knowledge to clinical trials with increased speed and accuracy^10,11^. Statistical models^12–16^ for data integration enhance the predictive power of gene-disease association, by incorporating prior knowledge of regulatory relation among different omics data and analysing them under one statistical framework.

With access to enormous data from several consortiums on various omics data, many data integration techniques have been developed so far. Such methods comprise of dimension reduction techniques, feature selection techniques using supervised, unsupervised or semi-supervised learning, graph or kernel-based techniques, Bayesian, and frequentist approaches^16–21^. But more often, integration methods combine multiple omics data from large consortiums pertaining to different cohorts^15,22^. Such methods are prone to spurious prioritisation of associated genes owing to substantial cross-cell-type variation^23^. For these reasons and to reduce the stratification bias due to population diversity, increasing attempts are being made to create large scale multi-omics datasets recently by combining multiple assays from the same set of samples^24^.

However, individual research groups made substantial efforts for generating data on genetic variation, gene expression, methylation, phenotype, etc. simultaneously from the same set of samples to study biomarkers. Realising its great potential, data sharing platforms/repositories stored such heterogeneous data for the broader markets immediate study^24–26^. But unlike large consortium data, these data might have a relatively small sample size. In addition, missing data occur across multiple omics. For example, while integrating information from different types of genomic data, typically, genotype information are available for all the individuals but gene expression and/or methylation information are often partially missing^27^. Yet gene expression and/or methylation assays are rarely repeated for generating the missing data due to various reasons such as the huge cost of the assays^2^, degradation of mRNA^28^ and/or dearth of tissue samples, etc. So, a major challenge is to integrate multi-omics data in presence of partially missing individual-level observations^24^.

Few Bayesian methods^29,30^ consider missing value imputation in multi-omics data integration model. But sometimes the imputed data overshadow the contribution from the partially observed data for certain percentages of missing data and generally involves a huge computational cost to decide whether or not to impute^29^. Moreover, imputing the missing values might be misleading^23^as it introduces bias and uncertainty in the data^31^, especially when the missing percentage is large and/or the reason of missing is unclear. To deal with the missing values it is important to understand the data source, data structure, missing mechanism, and amount of missing data ^32^along with its relation with the phenotype^31^. Some network-based methods^33^ considers partial multi-omics data integration using similarity network but assumes the same contribution of different omics.

In this paper, we propose a multi-omics genetic association tool, called TiMEG (Tool for integrating Methylation, gene Expression and Genotype), for identification of disease-associated biomarkers, by integrating single nucleotide polymorphism (SNP), gene expression, and DNA methylation with partially missing omics data under case-control paradigm. Our method elucidates the effect of multiple omics on the qualitative phenotype (case-control status). It jointly dissects the information on various omics data, their inter-relationship, and the information from individuals with completely as well as partially available omics data, without imputing the missing data. Using a likelihood-based approach, TiMEG models the conditional distribution of the response variable using the missing predictor variables ^34,35^.

The asymptotic distribution of our test statistic under the null hypothesis of no genetic association computes p-values much faster compared to other computational methods like permutation-based techniques etc. Extensive simulation confirms robust performance in terms of prediction accuracy of estimation in 10-fold cross-validation, controlled type I error rate, high statistical power, and consistency of the test under different missing data schemes. Application of our method on a real dataset of tuberous sclerosis (also called tuberous sclerosis complex (TSC)) patients and healthy controls (phs001357.v1.p1) identified functionally relevant genes and gene clusters belonging to the pathways that are involved in TSC pathogenesis. Even with a small sample size and a substantially high percentage of missing data, TiMEG could be used for the identification of biomarkers without losing any information. This leads to capturing weaker signals that remain unidentified in larger single-omics data analysis.

## 3 Results

### 3.1 TiMEG method

In presence of a limited sample size, missing individual-level information on multiple assays poses a great loss of information. Imputation might lead to bias in such a small sample size as the percentage of missing data is large. We introduce TiMEG, a general analytical approach for the identification of biomarkers associated with a disease by integrating multiple omics data with/without missing individual-level omics information under a case-control paradigm. We integrate data from DNA sequencing, gene expression, and DNA methylation assays along with covariates and qualitative phenotype (disease status) from the same set of samples. Figure 1 gives a general structure of omics data availability. Based on Figure 1, we design our missing data schemes (Table 1).

**Table 1:** Missing data schemes

| Data type<br>Sample size | Phenotype | Covariates | Genotype | Gene expression | Methylation |
| --- | --- | --- | --- | --- | --- |
| $n_1$ | ✓ | ✓ | ✓ | ✓ | ✓ |
| $n_2$ | ✓ | ✓ | ✓ | ✗ | ✓ |
| $n_3$ | ✓ | ✓ | ✓ | ✓ | ✗ |
| $n_4$ | ✓ | ✓ | ✓ | ✗ | ✗ |

Notation: ‘✓’ indicates ‘available’, ‘✗’ means ‘missing’ data. For $n_1$ individuals, no data is missing; for $n_2(n_3)$ individuals only gene expression (methylation) data are missing; both gene expression and methylation data are missing for $n_4$ individuals

**Figure 1:**
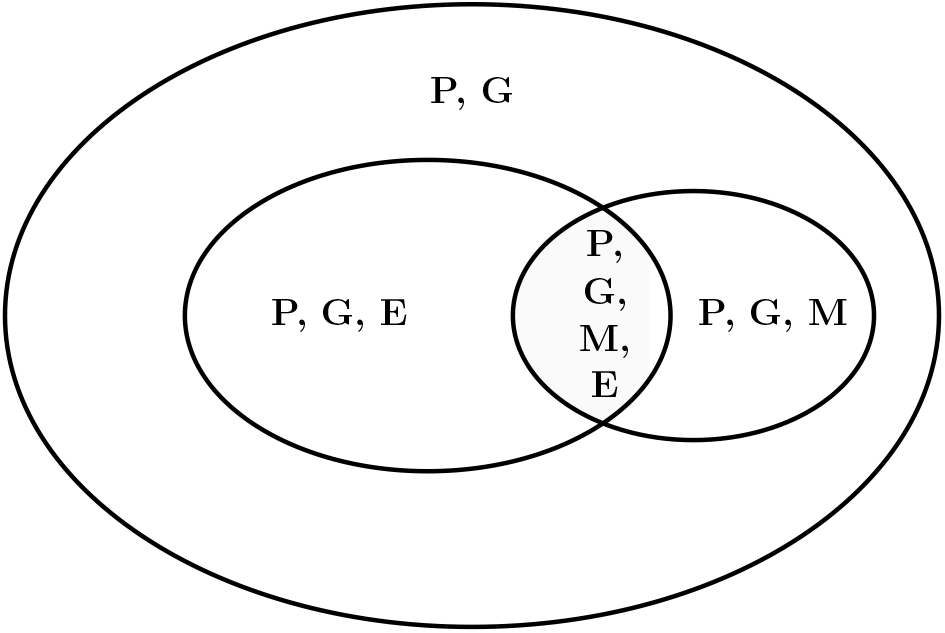
Structure of data availability for genotype (G), gene expression (E), methylation (M) and phenotype (P). Each letter indicates the presence of corresponding data

TiMEG relies on a likelihood approach to gather information on the missing data by estimating the parameters in the likelihood function containing incomplete omics data rather than imputing the missing data before the analysis. To obtain the likelihood function where one or more omics data are missing, we find the probability distribution of the response variable (disease status) conditional on the available omics information by (1) integrating out the missing variable from the joint likelihood function (see Methods), and (2) exploiting the interdependence among the multiple omics data.

TiMEG incorporates the inter-relationship among different omics data to provide additional information in the model. Since an individual’s gene expression level could be regulated by alteration in the DNA sequence, TiMEG considers the effect of genotype and methylation on gene expression. Similarly, it incorporates the effect of genotype on methylation and also the effect of genotype, methylation, gene expression, and covariates on disease status. We assume that after appropriate transformation and normalization procedures, individual-level gene expression and methylation data follows a bivariate normal distribution^13^.

### 3.2 Features of TiMEG

TiMEG is a statistical tool for the identification of disease-associated combinations of multiple omics. For example, a significant combination provides a gene along with a cis-methylation and a cis-genotype (collectively called a ‘trio’ throughout the paper). In this article, we illustrate a detailed pipeline for the identification of significant trios, if at least one component of the trio is associated with the disease.

Other advantages of TiMEG include the ability:

- to theoretically handle an unrestricted number of missing transcriptomic and epigenetic data as elucidated in the Methods section and confirmed using simulations,
- to capture weaker signals that remain unidentified in a single-omics analysis by incorporating more information from the inter-relation among multiple omics data in the likelihood framework,
- of robust performance in terms of prediction accuracy of estimation in 10-fold cross-validation, controlled type I error rate, and high statistical power,
- of efficient incorporation of correlated omics information to decipher significant signals resulting in reduced false-negative rate, and
- of prompt calculation of p-values using the asymptotic distribution as opposed to computation-intensive permutation-based methods.

Since alteration in gene expression regulation is expected to alter phenotype more than any change in the DNA sequence, the performance of TiMEG reveals that the effect (in terms of statistical power) of a certain percentage of missing gene expression data is more than the same percentage of missing methylation data (see Simulations). Missing both omics information for a subset of individuals will lead to much more loss of information (and therefore statistical power) than missing gene expression or methylation data on any subset of equivalent size. Thus, TiMEG agrees with the biologically accepted notion. Regardless of the sample size and/or percentage of missing omics data, wet-lab researchers will be able to promptly identify significant biomarkers from their data by calculating p-values using this tool.

### 3.3 Performance of TiMEG

#### 3.3.1 Simulations

We perform extensive simulations to study the performance of TiMEG for varying percentages of missing omics data under different missing data schemes. As more often gene expression and methylation data are missing for a subset of the genotyped individuals, we assume that genotype, phenotype, and covariate data are available for the entire sample of size *n* (say). As shown in Table 1, for these *n* individuals, there could arise four different scenarios (1) none of the other two omics data are missing for a certain subset of size *n*_1_(say), (2) only gene expression data are missing for another subset of size *n*_2_(say), (3) only methylation data are missing for a third subset of size *n*_3_(say), and (4) both of the omics data are missing for the remaining subset *n*_4_(say). Here, we consider two covariates, age and gender and simulate them from *N*(40, 6) and *Bin*(1, 0.5) respectively. First, we simulate data under scheme 1 i.e when there is no missing observation.

For generating genotype data, we assume an SNP having two alleles *A* and *a* with *A* as a minor allele. Considering di-allelic loci, we simulate genotype data from Bernoulli distribution assuming Hardy-Weinberg Equilibrium (HWE) for controls with minor allele frequency (MAF) 0.2 for associated SNP. We generate genotypes for cases using additive model for relative risk^36^ based on disease prevalence = 0.1, genotypes (AA, Aa, and aa), and relative risk = 1.2.

Next, we generate methylation and gene expression values using Eqs. 2 and 3 (see Methods) and assuming the values of the parameters as *α*_0_ = 1.3, *α_g_* = 2.4, *γ*_0_ = 1.9, *γ_g_* = 0.6, = 2.3. Methylation-gene expression pair follows a bivariate normal distribution with variances 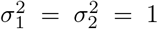. We assume that the means of the bivariate distributions for cases and controls differ by 0.3. Now, based on covariates, genotype, gene expression, and methylation, we simulate the phenotype of each individual using Bernoulli distribution in Eq. 1 (see Methods) with parameters *β*_0_ = 1, 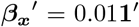, *β_g_* = 0.1, *β_m_* = 0.2, *β_e_* = 0.3. We generate data for *n* cases and *n* controls where the sample size n is taken as 100, 150 and 200. For the sake of power comparison, we take equal sample sizes for cases and controls. However, our method works for unequal sample sizes for cases and controls as well.

For the other three schemes, we generate complete data as above and remove some omics information to introduce missingness. For the second scheme where only gene expression is missing, we remove varying percentages (10%, 20%, 40%, 60%, 80%) of gene expression values. Similarly, we remove varying percentages of methylation values for the third scheme. For both omics missing scheme, we remove gene expression and methylation values for different combinations of missing percentages (Table 2 and 3).

**Table 2:** Type I error rate under different combination of sample sizes and varying percentages of missing methylation and/or gene expression values based on 10000 simulations

| $\begin{matrix} SS \\ \text{missing \%} \end{matrix}$ | 100 | 150 | 200 | $\begin{matrix} SS \\ \text{missing \%} \end{matrix}$ | 100 | 150 | 200 |
| --- | --- | --- | --- | --- | --- | --- | --- |
| (0,0,0) | 0.0534 | 0.0528 | 0.0533 | (10,0,10) | 0.0860 | 0.0847 | 0.0708 |
| (0,0,10) | 0.0836 | 0.0805 | 0.0757 | (0,10,10) | 0.0544 | 0.0519 | 0.0469 |
| (0,0,20) | 0.0828 | 0.0760 | 0.0758 | (10,10,0) | 0.0536 | 0.0493 | 0.0537 |
| (0,0,40) | 0.0758 | 0.0803 | 0.0776 | (20,0,20) | 0.0826 | 0.0810 | 0.0793 |
| (0,0,60) | 0.0826 | 0.0789 | 0.0741 | (0,20,20) | 0.0557 | 0.0500 | 0.0476 |
| (0,0,80) | 0.0869 | 0.0848 | 0.0761 | (20,20,0) | 0.0588 | 0.0502 | 0.0489 |
| (0,10,0) | 0.0552 | 0.0508 | 0.0497 | (40,0,40) | 0.0884 | 0.0773 | 0.0794 |
| (0,20,0) | 0.0538 | 0.0497 | 0.0520 | (0,40,40) | 0.0546 | 0.0507 | 0.0455 |
| (0,40,0) | 0.0552 | 0.0533 | 0.0516 | (10,10,10) | 0.0551 | 0.0551 | 0.0566 |
| (0,60,0) | 0.0558 | 0.0537 | 0.0507 | (10,20,10) | 0.0557 | 0.0527 | 0.0530 |
| (0,80,0) | 0.0641 | 0.0570 | 0.0507 | (10,10,20) | 0.0556 | 0.0512 | 0.0502 |
| (10,0,0) | 0.0869 | 0.0755 | 0.0747 | (20,10,10) | 0.0536 | 0.0574 | 0.0495 |
| (20,0,0) | 0.0866 | 0.0796 | 0.0760 | (20,10,20) | 0.0556 | 0.0511 | 0.0573 |
| (40,0,0) | 0.0813 | 0.0783 | 0.0796 | (20,20,10) | 0.0523 | 0.0497 | 0.0501 |
| (60,0,0) | 0.0836 | 0.0871 | 0.0766 | (10,20,20) | 0.0523 | 0.0515 | 0.0467 |
| (80,0,0) | 0.0982 | 0.0854 | 0.0836 | (20,20,20) | 0.0538 | 0.0510 | 0.0518 |

$\text{missing\% } (m_1, m_2, m_3) \equiv (m_1\% \text{ both missing, } m_2\% \text{ only methylation missing, } m_3\% \text{ only gene expression missing})$ ; $SS$ : sample size for case (or control)

**Table 3:** Power under different combination of sample sizes and varying percentages of missing methylation and/or gene expresssion values based on 1000 simulations

| $\begin{matrix} SS \\ \text{missing \%} \end{matrix}$ | 100 | 150 | 200 | $\begin{matrix} SS \\ \text{missing \%} \end{matrix}$ | 100 | 150 | 200 |
| --- | --- | --- | --- | --- | --- | --- | --- |
| (0,0,0) | 0.697 | 0.878 | 0.950 | (10,0,10) | 0.564 | 0.748 | 0.907 |
| (0,0,10) | 0.640 | 0.800 | 0.933 | (0,10,10) | 0.617 | 0.831 | 0.923 |
| (0,0,20) | 0.579 | 0.760 | 0.917 | (10,10,0) | 0.598 | 0.807 | 0.907 |
| (0,0,40) | 0.552 | 0.730 | 0.866 | (20,0,20) | 0.483 | 0.701 | 0.834 |
| (0,0,60) | 0.485 | 0.676 | 0.829 | (0,20,20) | 0.581 | 0.767 | 0.901 |
| (0,0,80) | 0.445 | 0.634 | 0.795 | (20,20,0) | 0.506 | 0.722 | 0.844 |
| (0,10,0) | 0.605 | 0.842 | 0.937 | (40,0,40) | 0.327 | 0.488 | 0.642 |
| (0,20,0) | 0.646 | 0.830 | 0.933 | (0,40,40) | 0.514 | 0.745 | 0.879 |
| (0,40,0) | 0.647 | 0.817 | 0.936 | (10,10,10) | 0.567 | 0.762 | 0.889 |
| (0,60,0) | 0.619 | 0.818 | 0.931 | (10,20,10) | 0.568 | 0.753 | 0.907 |
| (0,80,0) | 0.596 | 0.802 | 0.916 | (10,10,20) | 0.543 | 0.724 | 0.867 |
| (10,0,0) | 0.612 | 0.804 | 0.911 | (20,10,10) | 0.524 | 0.755 | 0.858 |
| (20,0,0) | 0.539 | 0.778 | 0.882 | (20,10,20) | 0.484 | 0.710 | 0.797 |
| (40,0,0) | 0.425 | 0.641 | 0.793 | (20,20,10) | 0.518 | 0.740 | 0.871 |
| (60,0,0) | 0.362 | 0.514 | 0.663 | (10,20,20) | 0.528 | 0.740 | 0.886 |
| (80,0,0) | 0.228 | 0.366 | 0.488 | (20,20,20) | 0.497 | 0.695 | 0.831 |

$\text{missing\% } (m_1, m_2, m_3) \equiv (m_1\% \text{ both missing, } m_2\% \text{ only methylation missing, } m_3\% \text{ only gene expression missing})$ ; $SS$ : sample size for case (or control)

To estimate the parameters in the model, we maximise the likelihood function using a numerical optimisation technique (see Methods). We construct a test statistic using the above estimates to test whether a trio is associated with the phenotype. Here, we use the likelihood ratio test for testing the null hypothesis (*H*_0_) of no effect of genotype, gene expression, and methylation on affection status. The asymptotic distribution of this test statistic follows a *χ*^2^ distribution with 3 degrees of freedom under *H*_0_. Figure 2 illustrates the QQ plot of sample quantiles from the empirical distribution of the test statistic under *H*_0_ with respect to theoretical quantiles of 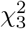 distribution, for a complete data and another dataset with missing data.

**Figure 2:**
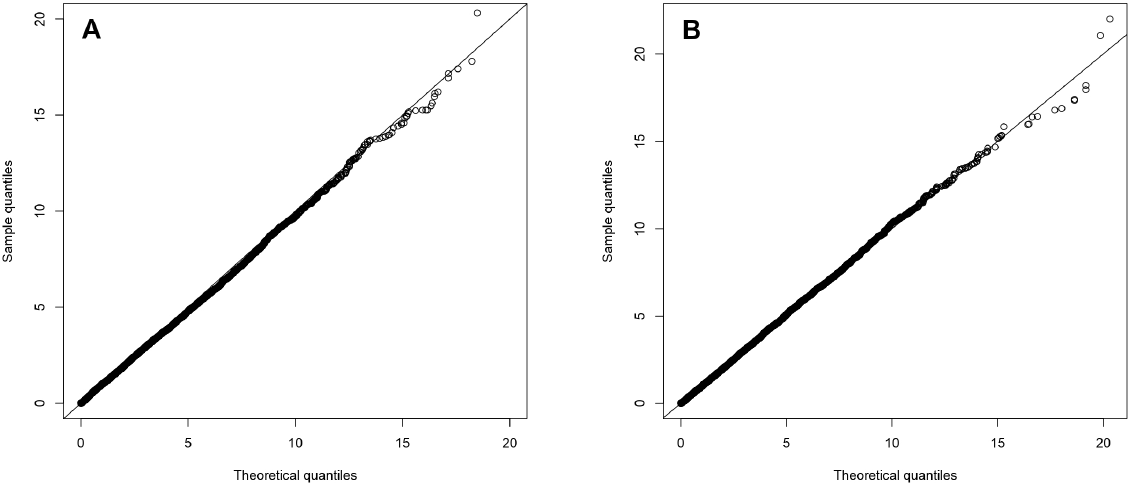
QQ-plot with sample size 200 based on the performance of simulated data. Figure 2A: QQ-plot with no missing data, Figure 2B: QQ-plot with 10% both gene expression and methylation missing, 10% only methylation and 20% only gene expression missing

Based on 10000 datasets, we find that type I error rate of our test is controlled nearly at 5% level of significance for each of the different sample sizes, missing data schemes, and percentages of missing omics data (Table 2). Hence, our test statistic is conservative in controlling false positives and is useful for p-value computation in a real dataset. To examine the performance of the test, we calculate statistical power under different missing data schemes and for various percentages of missing omics data based on 1000 datasets. To find whether the power of the test increases with an increase in sample size, we calculate power based on 5% cut-off points from 10000 datasets generated under *H*_0_, for different percentages of missing omics data corresponding to each missing data scheme. This would keep the type I error rate fixed exactly at 5% level so as to make a uniform power comparison. Table 3 demonstrates a substantial increase in power for every combination of missing data with increase in sample size. Under each missing data scheme, when the percentage of missing data increases, the power decreases. Evidently, when there is no missing data the power would be maximum. Thus, our test is consistent.

As mentioned earlier (in section 3.2), we now observe from Table 3, that TiMEG is more effected (as evident from the drop in statistical power) by (1) a certain percentage of missing gene expression data than the same percentage of missing methylation data and (2) missing both omics information for a subset of individuals than missing gene expression or methylation data on any subset of equivalent size. For instance, let us consider a fixed sample size 200 (say) and a fixed percentage of missing omics data 40% (say). From Table 3 we note that the power of missing only gene expression data (0.866) is less than that of missing only methylation data (0.936). Clearly, the power of missing both omics (0.793) is less than the minimum of the above two. Such a difference in power is biologically expected because alteration in gene expression is more informative than any other change in the DNA sequence. Thus, for a subset of individuals, no information on any of the two mentioned omics causes more loss of information compared to the presence of at least one of them. So, at the individual level, if possible, it is better to collect at least one observation from gene expression or methylation. Thus, provided there exists a choice, less percentage of missing gene expression is preferred than that of methylation because gene expression data is more informative.

When miscellaneous percentages of data are missing, we observe (from Table 3) a similar phenomenon as above. For the fixed overall percentage of missing omics data (40%) and the fixed sample size (200), we consider three combinations such as (1) 20% individuals have both omics missing, another 10% individuals have only gene expression missing, and another 10% individuals have only methylation missing, (2) 10% individuals have both omics missing, another 20% individuals have only gene expression missing, and another 10% individuals have only methylation missing, and (3) 10% individuals have both omics missing, another 10% individuals have only gene expression missing, and another 20% individuals have only methylation missing. The powers in the above three combinations are 0.858, 0.867, and 0.907. So we observe that, even in a miscellaneous missing scenario with such marginal difference in missing percentage of the omics, TiMEG is able to differentiate between statistical powers in accordance with the biological expectation. These results indicate that our method is able to capture all available information corresponding to every individual under study and its performance is robust to the percentage and scheme of missing omics data.

#### 3.3.2 Performance evaluation

To evaluate the predictive performance of our method, we assess the prediction accuracy of our estimation by 10-fold cross-validation (CV). We first generate a dataset that has a pre-assigned missing omics data structure and divide it into two parts, test set and training set. Observations with no missing data are then split into 10 blocks. One block is selected as a test set and all remaining individuals form a training set. Based on the first training set, we find an estimated *β* coefficients (using Eq. 1 in Methods) and classify the individuals in the corresponding test set to cases and controls. We repeat this procedure for all 10 test sets and calculate average prediction accuracy, specificity, and sensitivity for the dataset. For each pre-assigned missing omics data structure, we generate 100 datasets and perform 10-fold CV on each of them. We find that the mean prediction accuracy for classifying an individual to case or control group is more than 80% under all missing omics schemes and for different missing percentages (Table 4).

**Table 4:** Mean (±standard deviation) of prediction accuracy, specificity, sensitivity based on 10-fold cross validation of 100 datasets for each pre-assigned missing omics data structure

| missing % | Prediction accuracy | Specificity | Sensitivity |
| --- | --- | --- | --- |
| (0,0,0) | 83.275 ($\pm 1.954$) | 0.836 ($\pm 0.020$) | 0.822 ($\pm 0.024$) |
| (0,0,10) | 83.464 ($\pm 1.839$) | 0.835 ($\pm 0.022$) | 0.834 ($\pm 0.022$) |
| (0,0,20) | 83.406 ($\pm 1.993$) | 0.833 ($\pm 0.024$) | 0.833 ($\pm 0.024$) |
| (0,0,40) | 83.167 ($\pm 2.738$) | 0.825 ($\pm 0.034$) | 0.839 ($\pm 0.030$) |
| (0,0,60) | 82.775 ($\pm 3.040$) | 0.815 ($\pm 0.042$) | 0.841 ($\pm 0.039$) |
| (0,0,80) | 82.537 ($\pm 4.429$) | 0.815 ($\pm 0.059$) | 0.835 ($\pm 0.057$) |
| (0,10,0) | 83.531 ($\pm 1.773$) | 0.833 ($\pm 0.021$) | 0.838 ($\pm 0.022$) |
| (0,20,0) | 83.109 ($\pm 2.204$) | 0.827 ($\pm 0.027$) | 0.835 ($\pm 0.024$) |
| (0,40,0) | 83.133 ($\pm 2.259$) | 0.820 ($\pm 0.030$) | 0.843 ($\pm 0.026$) |
| (0,60,0) | 81.944 ($\pm 3.339$) | 0.805 ($\pm 0.041$) | 0.833 ($\pm 0.039$) |
| (0,80,0) | 82.500 ($\pm 4.533$) | 0.813 ($\pm 0.059$) | 0.837 ($\pm 0.058$) |
| (10,0,0) | 83.272 ($\pm 2.274$) | 0.832 ($\pm 0.026$) | 0.833 ($\pm 0.025$) |
| (20,0,0) | 83.087 ($\pm 1.970$) | 0.827 ($\pm 0.023$) | 0.834 ($\pm 0.021$) |
| (40,0,0) | 82.800 ($\pm 2.546$) | 0.820 ($\pm 0.031$) | 0.836 ($\pm 0.029$) |
| (60,0,0) | 82.687 ($\pm 3.070$) | 0.809 ($\pm 0.039$) | 0.844 ($\pm 0.034$) |
| (80,0,0) | 82.712 ($\pm 4.619$) | 0.808 ($\pm 0.058$) | 0.847 ($\pm 0.051$) |
| (5,0,5) | 83.092 ($\pm 1.892$) | 0.828 ($\pm 0.021$) | 0.833 ($\pm 0.022$) |
| (0,5,5) | 83.405 ($\pm 2.068$) | 0.831 ($\pm 0.023$) | 0.837 ($\pm 0.025$) |
| (5,5,0) | 82.967 ($\pm 1.828$) | 0.828 ($\pm 0.022$) | 0.831 ($\pm 0.021$) |
| (10,0,10) | 83.303 ($\pm 2.184$) | 0.829 ($\pm 0.027$) | 0.837 ($\pm 0.025$) |
| (10,10,0) | 82.991 ($\pm 2.134$) | 0.825 ($\pm 0.026$) | 0.835 ($\pm 0.025$) |
| (10,5,5) | 83.444 ($\pm 2.361$) | 0.831 ($\pm 0.028$) | 0.838 ($\pm 0.025$) |

missing% ( $m_1, m_2, m_3$ ) ≡ ( $m_1$ % both missing, $m_2$ % only methylation missing, $m_3$ % only expression missing)

We observe that median prediction accuracy remains the same under all scenarios except when the percentage of missing data is very high (Figure 3). Although the deviation of median prediction accuracy for an extremely high percentage (~ 80%) of missing values compared to that for no missing data is small, the higher dispersions indicate fluctuations of the prediction accuracies. This implies that the number of false-positive and false negative classifications fluctuates for these extreme missing scenarios. We illustrated this increase in dispersion through the plot of false-positive rate (1 - Specificity) versus misclassification rate ((1 – prediction accuracy)/100) under different percentages of missing omics data pertaining to different missing data schemes (Figure 4, S1-S3). It is evident that, for comparatively smaller percentages of missing data, the dispersions are much less. Only for extreme conditions of missing data, the dispersion is slightly higher. Thus, we find that our method provides a robust estimate of the parameters under different missing omics schemes with a reasonable missing percentage and has high predictive power.

**Figure 3:**
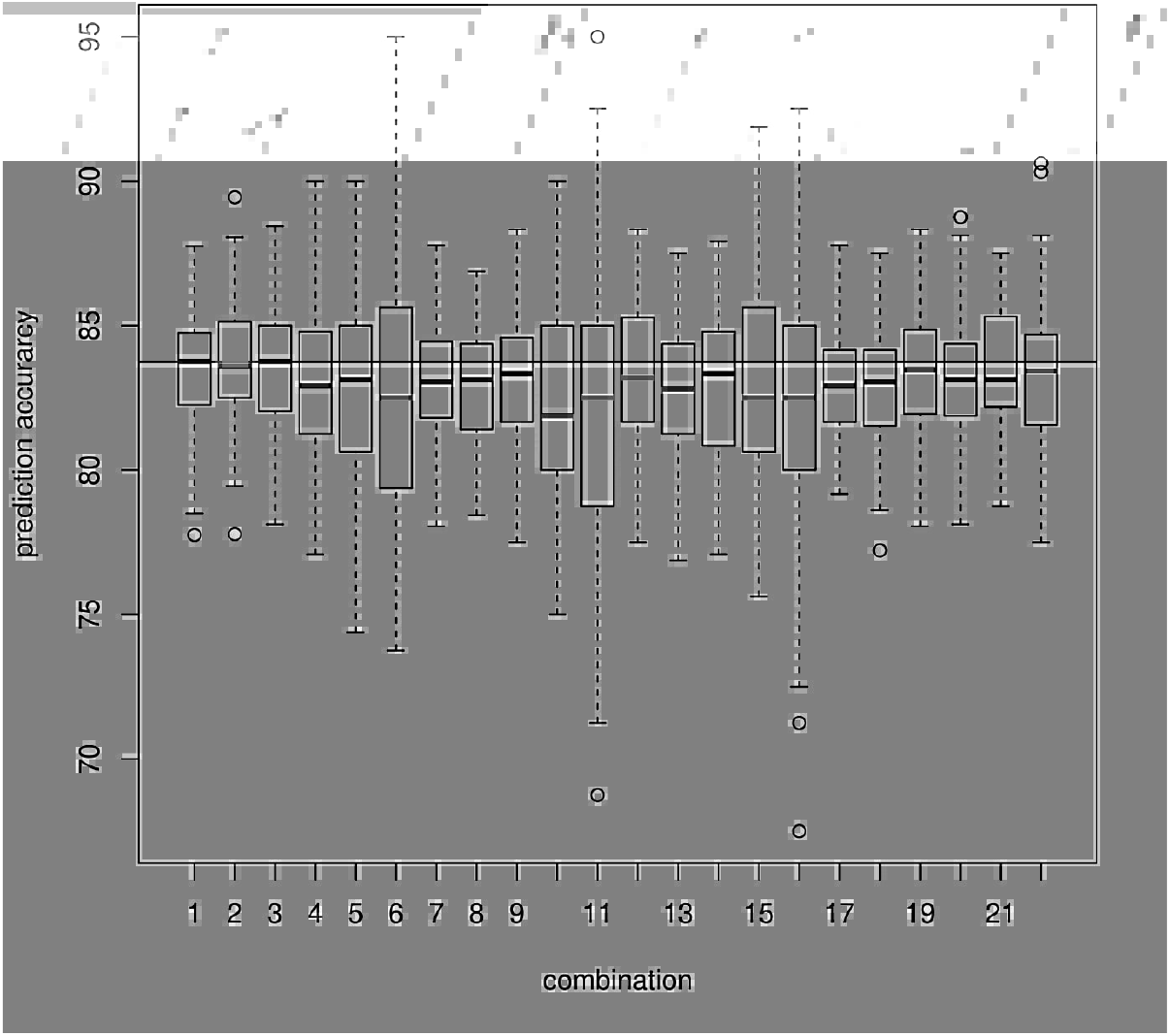
Boxplot of prediction accuracy from 10-fold CV based on 100 datasets each with 200 cases and 200 controls under different missing omics data structure. The black horizontal line indicates median prediction accuracy for the datasets with no missing information. Each boxplot (from left to right) signifies one combination each viz. no missing information, only 10%, 20%, 40%, 60%, 80% gene expression missing respectively, only 10%, 20%, 40%, 60%, 80% methylation missing respectively, 10%, 20%, 40%, 60%, 80% of both gene expression and methylation missing respectively, 5% of both missing along with 5% of only gene expression missing, 5% of only gene expression missing along with 5% of only methylation missing, 5% of both missing along with 5% of only methylation missing, 10% of both missing along with 10% of only gene expression missing, 10% of both missing along with 10% of only methylation missing, 10% of both missing along with 5% of only gene expression missing and another 5% of only methylation missing.

**Figure 4:**
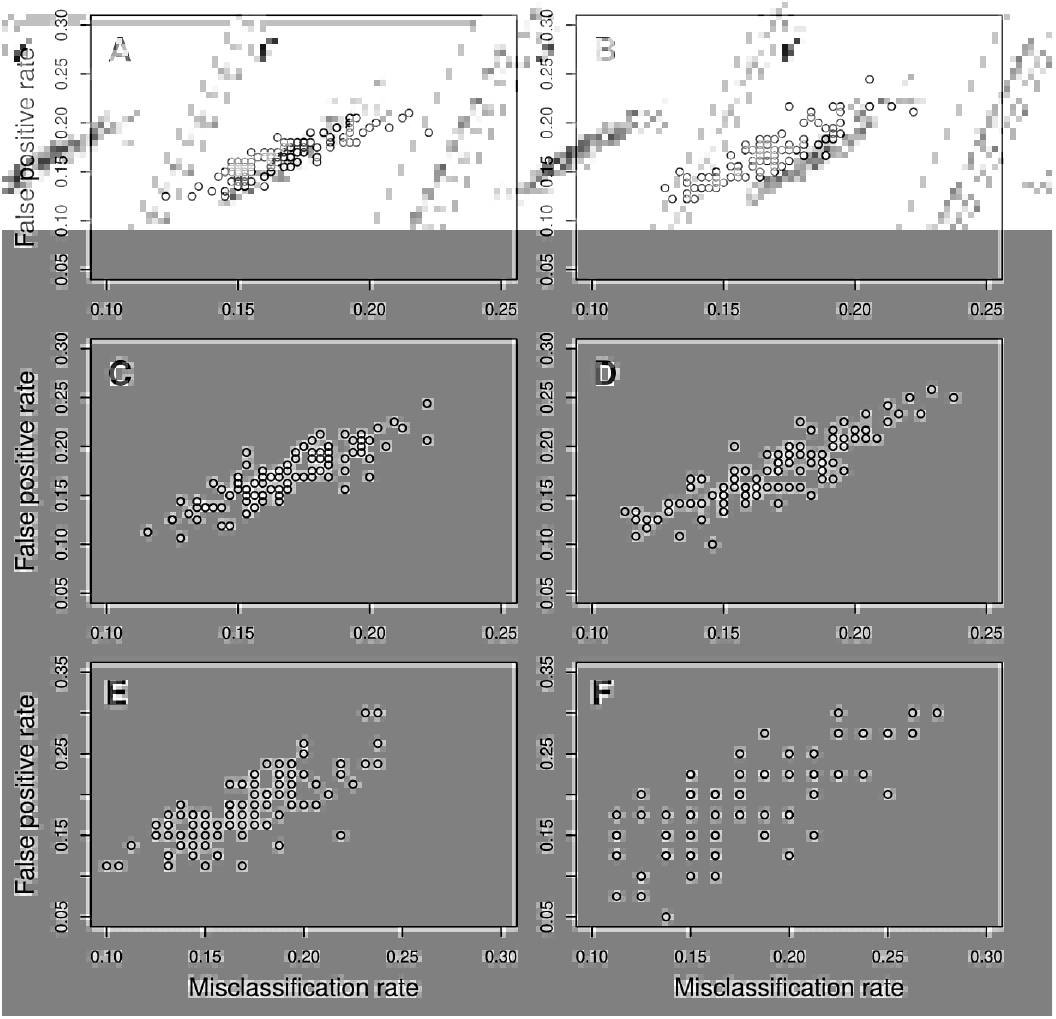
Plot of Misclassification rate vs False positive rate (1-Specificity) for only gene expression missing. Figure 4A depicts no missing data scenario while Figures 4B-4F respectively depict 10%, 20%, 40%, 60% and 80% only gene expression data missing scenarios

### 3.4 Application to a real dataset

We applied our proposed method to a dataset on Tuberous Sclerosis Complex (TSC) patients and healthy controls (phs001357.v1.p1)^27^ obtained from the database of Genotypes and Phenotypes (dbGaP). TSC is a rare genetic disorder that causes growth of non-cancerous (benign) tumours in the brain and other vital organs like kidneys, heart, skin, etc., and in some cases leads to significant health problems.

After processing raw data (see Methods), we obtained 8036 gene expression data on 27 cases and 7 controls, methylation data at 481470 CpG sites on 22 cases and 7 controls, and 1298477 whole-genome genotype data on 38 cases and 7 controls. We got data on all three omics for the control individuals. But only 12 case individuals had complete omics information. 9 other case individuals had all omics data except gene expression data, another group of 13 patients had no methylation data, and 4 patients had neither gene expression nor methylation data. Phenotype or disease status, covariates (such as age and gender), and genotype data were available for all cases and controls.

Since control samples are only a few, we considered those genes that have no missing gene expression value in controls, while in case samples we allow up to 50% missing gene expression value. Next, we find the SNPs that are within 2000 bp upstream and downstream of each gene. We considered these SNPs as the cis-SNPs to the gene. Moreover, if any methylation site is associated with a gene, information such as the corresponding gene name and chromosome number are known from dbGaP. So, methylation sites in the vicinity of a gene are considered as cis-CpG sites corresponding to the gene. After filtering the data, we find the number of unique genes containing at least one cis-SNP and one cis-CpG site reduces to 1691 and the total number of trios (comprised of one gene expression with one cis-genotype and one cis-CpG site corresponding to the gene) is 1184436. In order to identify the trios associated with the disease, we perform our test for all the mentioned trios, followed by Benjamini-Hochberg (BH) multiple corrections (across all tests).

#### 3.4.1 Interpretation of a TiMEG trio

For each significant trio, one or more of its components are associated with the disease. However, we are more interested to observe whether TiMEG is able to identify loci with moderately low effect sizes that are missed by single-omics analyses. Therefore, we find those combinations where TiMEG shows association but not the separate analyses. We see that TiMEG successfully captures weaker signals that remain unidentified in single-omics analyses. The probable reason might be that, single-locus from any omics data is unlikely to account for much of the variability in the phenotype. Moreover, it is often indicated that an increase in the sample size might capture the loci with moderate or low effect on the disease, but in that case, multiple testing burden also increases resulting in missing out true signals. But our method reduces false-negative associations by efficiently incorporating correlated omics information to decipher the significant signals associated with the disease. Particularly in this article, TiMEG tests if there is any effect of at least one of the components of the trio on the phenotype, but it is also able to test the effect of a single omics locus or combination of any two omics (see Methods). Emphatically, testing a single omics locus using TiMEG will provide greater insight than traditional single-omics analyses because of incorporating additional information from other available omics in the integrated model.

#### 3.4.2 Functional annotation of TSC genes

It is well known that mutation in either of the two tumor-suppressor genes viz. *TSC1*^37^ and *TSC2*^38^, that code for hamartin and tuberin proteins respectively are responsible for TSC. The hamartin/tuberin heterodimer encoded by the interaction of *TSC1* and *TSC2* gene products, function in complex pathways^39^. *TSC*1/2 genes and hence the hamartin/tuberin complex plays a fundamental role in the regulation of phosphoinositide 3-kinase (PI3K) signaling pathway^40^that inhibits the mammalian target of rapamycin (mTOR) through activation of the GTPase activity of Rheb ^41^.

Using TiMEG we obtained 170 unique genes (see Supplemental Table S1) from 3283 significant trios (https://github.com/sarmistha123/TiMEG), to be associated with the disease risk. These trios are significant due to the combined effect of all its components but none of the single-omics analyses could identify any of the corresponding components. Among the contents of this list, there exits a trio corresponding to gene *TSC1* with cis-genotype kgp7096367 and cis-methylation site *cg*19350728 that shows no association in any of the single-omics analysis but their combined effect is significant. TSC2 gene is excluded from our analysis because of a mismatch between probe id and HGNC IDs (See Methods).

We use David software^42,43^ to identify the functional annotations of these genes. Based on David’s group enrichment score, we obtained 5 clusters of our genes. The cluster with the maximum group enrichment score is associated with the serine/threonine kinase pathway. This pathway has a strong functional relation with TSC disease, as it is known that mutations in *TSC*1/2 genes impair the inhibitory function of the hamartin/tuberin complex, leading to phosphorylation (activation) of ribosomal protein S6 kinase beta-1 (S6K1), a serine/threonine kinase which is a downstream target of mTOR^41^.

Another cluster is associated with the zinc-finger protein pathway. Recent findings have highlighted the importance of the zinc-finger family and its involvement in tumorigenesis^44^. Interestingly, this pathway has some special implications in terms of brain tissues. Protein associated with Myc (called Pam) that is abundantly expressed in the brain, has been found to be associated with the tuberin/hamartin complex^45^. The C terminus of Pam containing the RING zinc-finger motif binds to tuberin^45^. Besides, Pam is a highly conserved nuclear protein that interacts directly with the transcriptional-activating domain of Myc (a protooncogene that plays an important role in the regulation of cellular proliferation, differentiation, and apoptosis and can contribute to tumorigenesis) ^46^and regulates mTOR signalling ^47^.

Studies have revealed that TSC receives inputs from at least three major signaling pathways (PI3K-Akt-mTOR, ERK1/2-RSK1, LKB1-AMPK) in the form of kinase-mediated phosphorylation events that regulate its function as a GTPase activating protein (GAP)^48^. But only two genes viz. *TSC1/2* are widely known to be responsible for the disease. Therefore, we searched whether any of our significant genes belonging to the same pathway as that of *TSC1* gene. Using David software we identified genes *ACACA* and *CREB5* in our list of significant genes, that occur in two pathways to which *TSC1* belongs.

Studies show that *TSC1* deficiency elevated *ACACA* expression and fatty acid synthesis, leading to impaired epigenetic imprinting on selective genes; tempering *ACACA* activity was able to divert cytosolic acetyl-CoA for histone acetylation and restore the gene expression program compromised by *TSC1* deficiency^49^. *CREB5* encodes CREB protein that serves as transcriptional activator of Rheb. Rheb acts as an immediate activator of mTOR and in turn promotes tumorigenesis independently of *TSC2*^50^. Moreover, we identified other genes such as *JAK3,GNG4,FGFR2,EFNA2,LAMC2* that belong to one of the pathways as that of *TSC1*.

#### 3.4.3 Implication of TiMEG

It is important to note that these genes could not have been identified by single-omics analyses. Data integration of different omics led to these findings even when the sample size is small and individual-level data is not available on all omics. Thus, TiMEG holds the potential to understand the genetic architecture of a disease etiology by combining three different omics data, even in presence of missing omics data, and when the sample size is not humongous.

Thus, if the sample size is not huge, studying single-omics data to find any new gene that is susceptible to the disease risk is difficult because most of the known diseases have already been studied extensively. Moreover, scientists nowadays are interested in developing drug targets with genetic evidence of disease association as they are much more likely to get approved ^51^. So, the identification of new disease-associated genes or biomarkers are immensely important to understand the relation of disease with various genes in the pathways. Downstream/detailed investigation of these biomarkers could provide a better understanding of the disease etiology and hence discovering best drug targets that might lead to successful development of novel drugs.

## 4 Discussion

Multi-omics data integration elucidates understanding the genetic architecture of diseases and complex traits, by incorporating additional information from different types of genomic data. But the presence of missing values poses a major challenge. This is more crucial when the sample size is limited and/or the percentage of missing data is large. Sometimes, these missing values occur due to biological reasons such as degradation of RNA or other technical issues. But when resources are limited, more often these assays are not repeated for the missing omics data. Typically, genotype data, being less expensive than gene expression and DNA methylation assays, are available for the entire sample. One option to analyse such data is using a sub-sample for which data are available for all omics. Clearly, such a complete case analysis loses a great deal of information. Again, imputation might induce bias arising due to the genetic diversity of reference data^15,22^. On the other hand, different types of omics data may be correlated and associated with a disease, directly or indirectly. Thus, integrating evidence from the inter-relationship among omics data provides additional information for biomarker identification.

We propose TiMEG, a tool for the identification of biomarkers integrating genotype, gene expression, and DNA methylation in presence of missing data under the case-control paradigm. Based on a likelihood approach, TiMEG is able to capture weaker signals that are often missed by single-omics analysis, by efficiently combining the information on interdependence among multiple omics data. Rather than imputing the missing data before the analysis, TiMEG accumulates information on the missing data by estimating the parameters in the likelihood function containing incomplete omics data. For calculating the likelihood function for incomplete data, we evaluate the conditional distribution of the response variable given the available information. This information not only includes the available omics data but also the inter-relationship among different omics. Moreover, our method has the ability to tackle an unrestricted number of missing transcriptomic and epigenetic data. Asymptotic distribution of our test statistic derived under the null hypothesis of no association will lead to the fast calculation of p-values compared to computation-intensive permutation-based techniques. Moreover, the normal approximation of the sigmoid function ^52^ reduces the computation time to a great extent. Thus, TiMEG could be promptly applied by the end-users on real datasets.

Simulation results confirm consistency of the test, robust performance in terms of prediction accuracy of estimation in 10-fold CV, controlled type I error rate, and high statistical power. Moreover, as the percentage of missing values increases, the power of the test decreases as expected. Our method shows such robust performance because we observe in the simulation that, for upto moderately high percentages of missing data, the power and 10-fold prediction accuracy of estimation are close to that of no missing data. Simulation results also indicate the reduction in power of the test is not substantial for extremely large missing percentages. Besides, the median prediction accuracy is nearly the same under all scenarios except when the percentage of missing data is very high (Figure 3) but, the mean prediction accuracy of classifying an individual to case or control group is nearly the same (Table 4). Only for extreme percentages of missing omics data, fluctuations in misclassification rate are slightly higher (Figure 4, S1-S3). Thus, one of the major advantages of TiMEG lies in its applicability to moderately large missing percentages and limited sample size for identification of biomarkers. Another advantage is that it identifies a combination of multiple omics loci (or trio) as biomarkers. So, even when any one or all the components of a significant trio has a small effect on the disease, TiMEG detects it. This is because, it is able to integrate multiple omics loci with small effects together, such that their combined effect on the disease is moderately large.

More often, wet-lab researchers encounter missing data in multiple omics assays, when data are collected on both patients and matched controls. One example of such an experiment is by Martin et al.^27^. We applied our method to their real dataset related to TSC that we obtained from dbGaP. The dbGaP data had genotype for all individuals (both TSC patients and healthy controls) but gene expression and/or methylation data were missing for a number of TSC patients. Although mutations in TSC1 and TSC2 genes are widely known to be responsible for the occurrence of TSC disease, several studies illustrated evidence of factors other than mutations in these genes, to be involved in the etiology of the disease. Our method could identify a few more TSC associated genes at a much smaller sample size combining different omics data. Some of the identified genes have been previously reported to actively participate in the TSC disease causation^50,53^.

Although TiMEG tests a trio for possible association with a disease, it could be extended to test multiple SNPs and multiple methylation sites along with gene expression. But, this will increase the number of parameters in the model. One possibility is to replace multiple SNPs and methylation values by some combined value or score, for example, the median of the methylation values under study etc. We plan to extend TiMEG for the accommodation of multiple SNPs and CpG sites as future work. We have not considered any interaction effect among different omics on the phenotype in this model. So, another extension of this work would be considering the interaction effect. Moreover, extending TiMEG to accommodate mutations instead of SNPs is important for experiments related to cancer. Some experiments collect data on quantitative phenotypes. TiMEG could be applied in such cases by dichotomising the quantitative phenotype but it would lose information. Therefore, it is important to develop a method for quantitative phenotypes, which might not be straightforward. However, another strength of TiMEG is that it can test the effect of a single omics locus or combination of any two omics immediately. Such tests of single omics locus would provide greater insight than traditional single-omics analyses due to the additional insights from the other omics data.

Moreover, application of TiMEG to the available data from public repositories might enhance understanding of disease by identification of different biomarkers. A detailed functional analysis of the significant association signals might facilitate understanding of the intricate genetic architecture of disease and therefore, translate the potential stored in the genomic data to develop targeted therapies and aid in precision medicine research.

## 5 Methods

### 5.1 Model

Figure 1 provides a general missing data structure of multiple omics in different studies. The effect of this structure on the identification of biomarkers is much more prominent in studies with a limited sample size compared to large consortium data. The objective of TiMEG is to identify disease-associated biomarkers by integrating individual-level information from genetic (SNP), transcriptomic, and epigenomic data along with their phenotype (disease status), and covariates in such scenario. So, TiMEG explores the effect of multi-omics data on disease status and the interrelation among multiple omics for biomarker identification. To illustrate the scenario, we consider n individuals with a known binary qualitative phenotype. For individual *i* (*i* = 1, 2,…, *n*), let *Y_i_, G_i_, M_i_* and *E_i_* denote respectively phenotype, genotype, methylation, and gene expression, and *X_i_* denote the vector of *J* covariates like age, gender, and other environmental variables. We denote *Y_i_* = −1, for controls and *Y_i_* = 1, for cases. Conventionally, *G_i_* takes value 0, 1, 2 depending on the number of minor alleles present. Let **M** and **E** denote the vectors of continuous values for *n* individuals and *X* be a matrix of order *n ×J* that includes covariate values for all *n* individuals. Based on the aforementioned omics data, we propose the following model for the *i^th^*(*i* = 1,…*n*) individual.

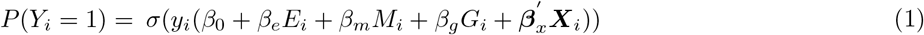

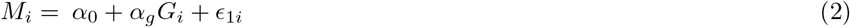

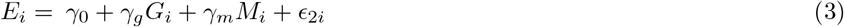

where 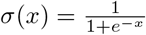. We assume that *G_i_ ~ Bin*(2, *p*) where, *p* is the probability of occurrence of a minor allele and 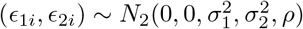 where 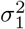 and 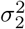 denote the variances of *ϵ*_1*i*_ and *ϵ*_2*i*_ respectively, and *ρ* = *Cor*(*ϵ*_1*i*_, *ϵ*_2*i*_). Here, we consider a likelihood based approach for estimation of the parameters in Eqs. 1 – 3. We denote the set of all parameters as, *θ* = (*β*_0_, *β_e_, β_m_, β_g_, β_x_, α*_0_, *α_g_, γ*_0_, *γ_g_, γ_m_, σ*^2^, 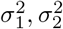, *p*)′. So, our joint likelihood function for the full data (i.e. when there is no missing observation), will be:

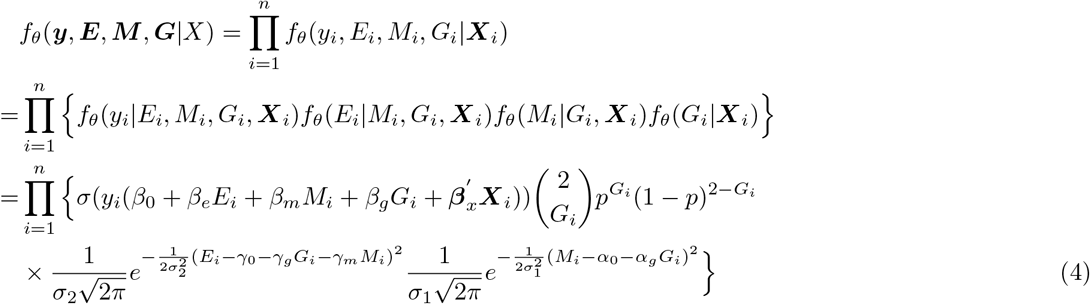

As discussed earlier, genetic variants are available for a large population but transcriptomic and epigenomic data tend to be missing due to various reasons. So, we assume that genotype, phenotype, and covariate data are available for the whole population while varying percentages of either gene expression or methylation or both the omics are missing (Table 1). In the following section, we introduce different schemes of missing values across multiple platforms.

### 5.2 Missing values scheme

We suppose that among *n* individuals, *n*_1_ individuals have complete data on all omics, phenotype, and covariates, for *n*_2_ individuals only gene expression is missing, *n*_3_ has only methylation values missing, and *n*_4_ individuals neither have data on gene expression nor on methylation. Thus, depending on the missing data type(s), we can consider three schemes of missingness (Table 1). In each case, we shall write the appropriate likelihood function. For that, we need to consider the following lemma (for proof see Appendix A, Supplemental Material).

#### Lemma 1.

*If φ*(*x*) *is the p.d.f. of a standard normal distribution, i.e*. 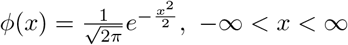, *then*

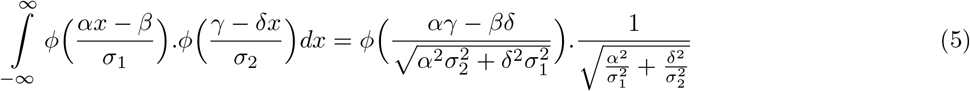

*where α, β, γ, δ*, 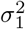 *and* 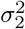 *are constants*.

Moreover, while deriving the likelihood functions, we approximate the logistic function in equation (4) by cumulative distribution function of a normal variable^52^. This approximation relation is given by:

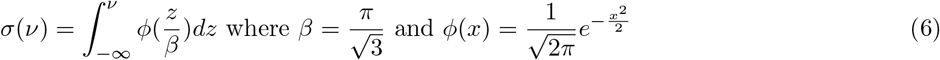

#### 5.2.1 Scheme 1: Only partial gene expression data are missing

Consider a situation where phenotype, genotype, covariate, and methylation data are available for all *n* individuals but, gene expression data are missing only for *n*_2_ individuals. This indicates that *n*_3_ = *n*_4_ = 0, and *n*_1_ = *n – n*_2_. Based on this missing observation scheme, we need to write the likelihood function using Lemma 1. But before that, we introduce a few notations for the sake of lucidity.

***Z**_i_* = (1, ***X**_i_, G_i_, M_i_, E_i_*)′, ***w*** = (*β*_0_, 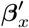, *β_g_, β_m_, β_e_*)′, ***Z**_i,o_* = (1, ***X**_i_, G_i_, M_i_*)′, ***Z**_i,m_ = E_i_, **w**_o_* = (*β*_0_, 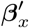, *β_g_, β_m_*)′. Now, in order to rewrite the likelihood function as in (4), we need a precise expression for *P*(*y_i_|**Z**_i_*), as given in the following result. Note that Result 1 (for Proof see Appendix B, Supplemental Material) is related to only n2 individuals for whom gene expression data are not available.

##### Result 1.

*Using the model (1-3)*,

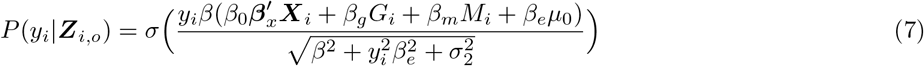

*where μ*_0_ = *γ*_0_ + *γ_g_G_i_ + γ_m_M_i_, for each i ∈ S_−E_, the set of n*_2_ *individuals for whom gene expression data are not available*.

Now without any loss of generality, we assume that for the first *n*_1_ individuals all data are available whereas the last *n*_2_ individuals do not have gene expression data. Hence, using Result 1, the likelihood function (4) can be written under the scheme 1 as:

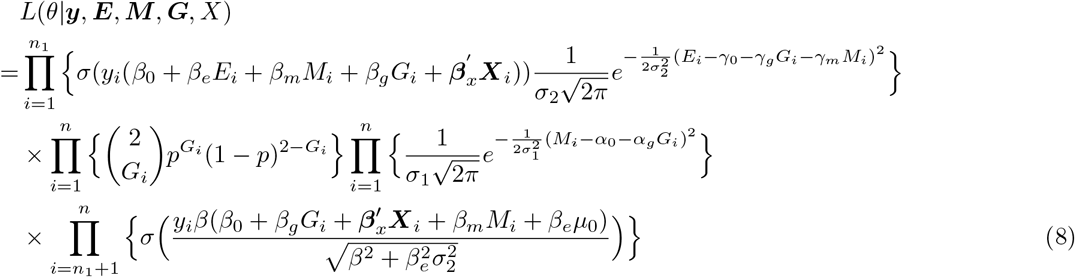

where *n*_2_ = *n − n*_1_.

#### 5.2.2 Scheme 2: Only partial methylation data are missing

We may have a situation where all types of data are available for *n*_1_ individuals and for another group of *n*_3_ individuals all types of data except methylation data are available. So here we have *n*_2_ = *n*_4_ = 0 and *n*_3_ = *n − n*_1_. So, the terms involving *M_i_* are not available for *n*_3_ individuals. Again as in the above scheme, we now introduce few notations as:

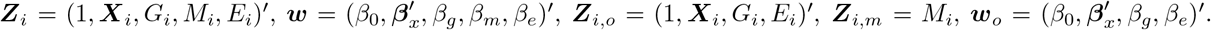

To write down the likelihood function, we first evaluate the expression for *P*(*y_i_|Z_i_*) as given in Result 2 using Lemma 1. Note that Result 2 (for proof, see Appendix B, Supplemental Material) is related to only n3 individuals for whom methylation data are not available.

##### Result 2.

*Using the model (1-3), for each i ∈ S_−M_*,

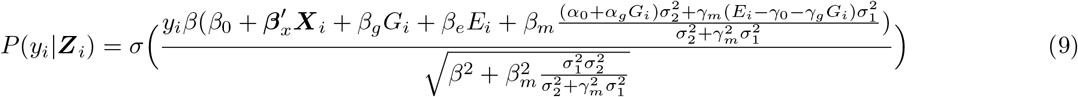

*where S_−M_ is the set of n*_3_ *individuals for whom no methylation data are available*.

Now without any loss of generality, we assume that for the first n_1_ individuals all data are available whereas the last *n*_3_ individuals do not have methylation data. Hence, using Result 2, the likelihood function (4) can be written under the scheme 2 as:

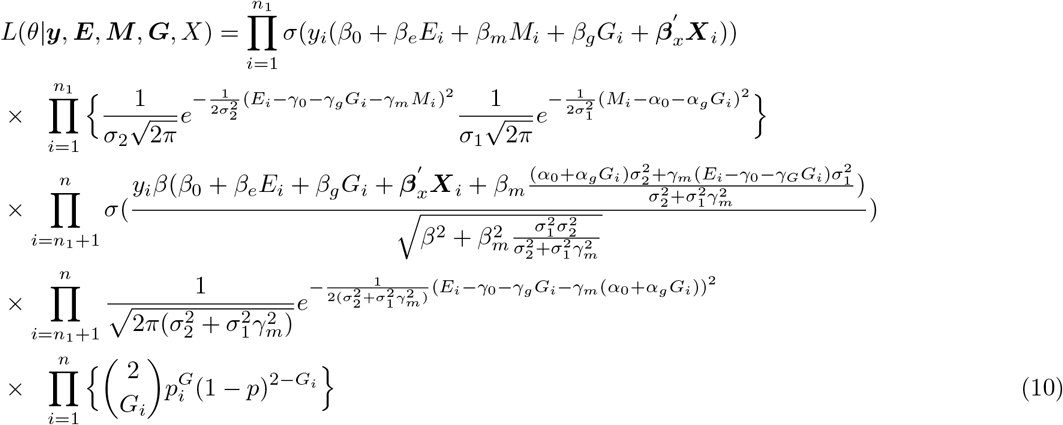

where, *n*_3_ = *n − n*_1_

#### 5.2.3 Scheme 3: Methylation and gene expression data are partially missing

Lastly, under the most general missing value scheme, we consider *n*_2_ individuals have only missing gene expression values, *n*_3_ individuals have only missing methylation values, *n*_4_ individuals have both missing gene expression and methylation values, and the rest of the individuals have all types of data. Similarly, as for other schemes, we now introduce few notations as:

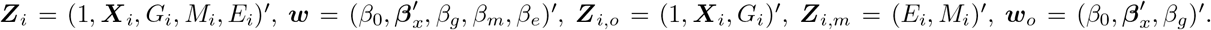

Then, we evaluate *P*(*y_i_|**Z**_i_*) as given in Result 3 (for proof, see Appendix B, Supplemental Material) using Lemma 1 in order to write the joint likelihood equation.

##### Result 3.

*Under the model (1-3), for each i ∈ S_−(E,M)_*,

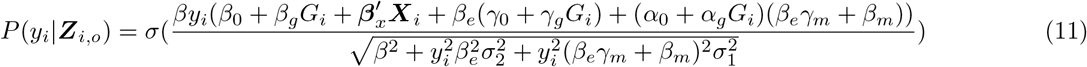

*where S_−(E,M)_ is the set of n*_4_ *individuals for whom both expression and methylation data are missing*.

In order to write down the likelihood function, we assume, without any loss of generality, that first *n*_1_ individuals have all data, next *n*_2_ individuals have all data except gene expression data, next *n*_3_ individuals have all data except methylation data and for the remaining *n*_4_ individuals neither gene expression data nor methylation data are available but phenotype, covariate, and genotype data are available. Clearly *n*_4_ = *n − n*_1_ − *n*_2_ − *n*_3_.

Using Results 1 – 3, the likelihood function (4) under scheme 3 can be written as:

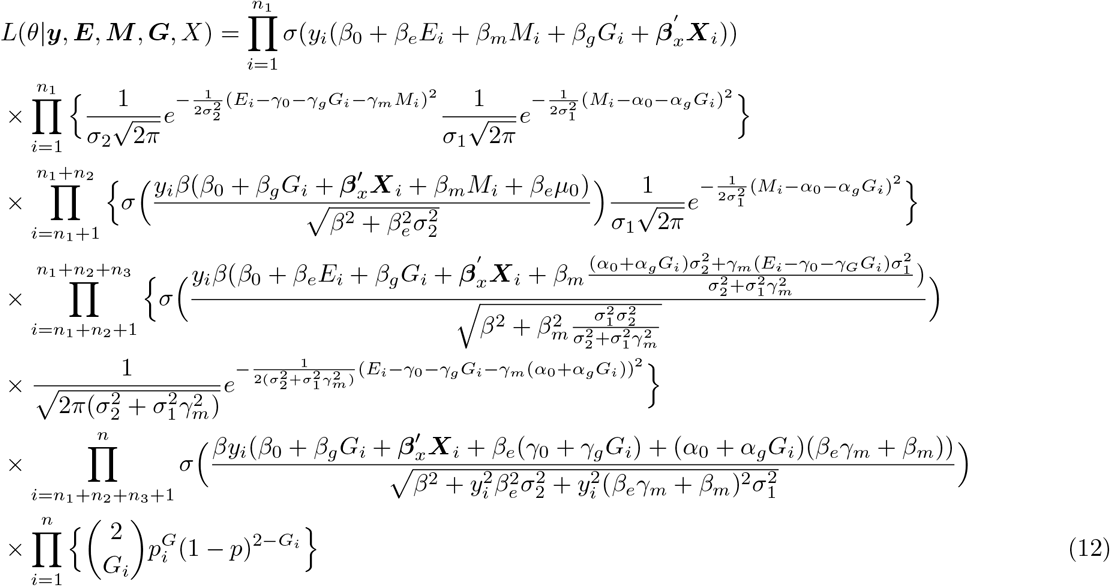

where 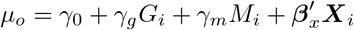, *n*_4_ = *n − n*_1_ − *n*_2_ − *n*_3_.

Next, for estimating the parameters in each of the likelihood functions, we use the L-BFGS-B method (in R package ‘stats’), an iterative algorithm for numerical optimisation to find the maximum likelihood estimates of the parameters. Thus, theoretically, our method is able to incorporate any amount of missing gene expression and methylation data. In this paper, we focus on the identification of disease-associated trios (that is, a combination of the gene along with its cis-genotype and cis-methylation site). Thus, the components of a significant trio is expected to have a joint effect on affection status. Traditional single-omics analyses of each component are likely to miss these loci unless the sample size is humongous and/or the technologies are tremendously improved. More information from multiple omics on each individual, coupled with additional insights from the inter-relationship among the omics, support the identification of significant loci even at smaller sample sizes compared to large single-omics analyses.

### 5.3 Hypothesis of interest

With the objective to identify a trio that may be associated with the disease or phenotype, we formulate the hypotheses of interest as:

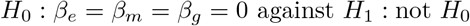

Rejection of *H*_0_ would indicate association with one or more components of the trio with the disease. To test the null hypothesis we adopt likelihood ratio test under a very general likelihood structure under various schemes of missing data as discussed in Section 3.3. The test statistic for testing *H*_0_ would be,

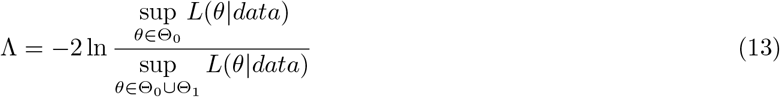

where *θ* is the vector of parameters in the likelihood function *L*, Θ_0_ and Θ_1_ are the parametric spaces under *H*_0_ and *H*_1_ respectively. Using standard asymptotic theory, it can be easily shown that the test statistic Λ follows *χ*^2^ distribution with 3 degrees of freedom asymptotically under *H*_0_. Usually, the sample sizes are considerably large so that we can use the asymptotic distribution of Λ under *H*_0_ for a real dataset. This reduces a huge computational burden while calculating the p-value in order to come to a conclusion.

If interested, one may test the effect of any two components (duos) such as genotype and gene expression, by testing

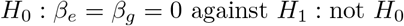

to find whether any combination of a gene and a genotype is associated with the disease. Other alternative hypotheses may be framed as per the objective. But here we consider only the identification of significant trios.

### 5.4 dbGaP data on TSC

We obtained a real data from dbGaP (phs001357.v1.p1)^27^ on TSC patients and healthy controls. Next, we analysed raw BAM files for gene expression data, IDAT files for genotype, and methylation data from brain tissues only. Genotypes are generated using Illumina Infinium Omni2.5 SNP arrays, methylation using Illumina Infinium HumanMethylation450 (HM450) BeadArrays, and gene expression using mRNA sequencing (RNAseq) for patient and control samples. For cases and controls, we derive log-normalised count for gene expression data, normalised-beta count for methylation data, and genotype data using Bioconductor package ‘DESeq2’, ‘methylumi’, and ‘CRLMM’ respectively in R software. For each probe ID, we find its transcription start and end sites according to the human genome assembly 19 (hg19) from the UCSC genome browser using Bioconductor package ‘TxDb.Hsapiens.UCSC.hg19.knownGene’ (Carlson and Maintainer 2015). To have the same gene nomenclature across all omics platforms we convert probe IDs to gene names (or HGNC IDs) (using http://hgdownload.cse.ucsc.edu/goldenPath/hg19/database/refFlat.txt.gz) and annotate the SNPs using Bionconductor package ‘humanomni258v1aCrlmm in R’.

## Acknowledgement

We are grateful to the Department of Biotechnology, Govt. of India, for their partial support to this study through SyMeC. SD thankfully acknowledges the support from the Department of Science and Technology, Government of India (SR/WOS-A/PM-1014/2015(G)).

## Data access

The TSC dataset analysed in this study is available from the database of Genotypes and Phenotypes (dbGaP study accession: phs001357.v1.p1).

## Code availability

The code for using TiMEG pipeline is freely available at https://github.com/sarmistha123/TiMEG. We also provide a small dataset as an example and explicit instructions for using this tool.

## Notes

https://www.ncbi.nlm.nih.gov/projects/gap/cgi-bin/study.cgi?study_id=phs001357.v1.p1

### Competing Interest Statement

The authors have declared no competing interest.

