## Supplemental Material for "TiMEG: an integrative approach for partially missing multi-omics data with an application to tuberous sclerosis"

Sarmistha Das,[1] and Indranil Mukhopadhyay[1,*]
[1]Human Genetics Unit, Indian Statistical Institute, Kolkata, West Bengal, 700108, India

### Appendix A

**Lemma 1.** *If $\phi(x)$ is the p.d.f. of a standard normal distribution, i.e. $\phi(x) = \frac{1}{\sqrt{2\pi}}e^{-\frac{x^2}{2}}$, $-\infty < x < \infty$, then*

$$\int\limits_{-\infty}^{\infty} \phi\Big(\frac{\alpha x-\beta}{\sigma_1}\Big).\phi\Big(\frac{\gamma-\delta x}{\sigma_2}\Big)dx = \phi\Big(\frac{\alpha\gamma-\beta\delta}{\sqrt{\alpha^2\sigma_2^2+\delta^2\sigma_1^2}}\Big).\frac{1}{\sqrt{\frac{\alpha^2}{\sigma_1^2}+\frac{\delta^2}{\sigma_2^2}}} \tag{1}$$

*where $\alpha$, $\beta$, $\gamma$, $\delta$, $\sigma_1^2$ and $\sigma_2^2$ are constants.*

*Proof.*

$$\phi\Big(\frac{\alpha x-\beta}{\sigma_1}\Big).\phi\Big(\frac{\gamma-\delta x}{\sigma_2}\Big) = \frac{1}{2\pi}\exp\Big\{-\frac{1}{2}\Big\{\frac{(\alpha x-\beta)^2}{\sigma_1^2}+\frac{(\gamma-\delta x)^2}{\sigma_2^2}\Big\}\Big\}$$

$$=\frac{1}{2\pi}\exp\Big\{-\frac{1}{2}\Big\{\Big(\sqrt{\frac{\alpha^2}{\sigma 1^2}+\frac{\delta^2}{\sigma_2^2}}\Big(x-\frac{\alpha\beta\sigma_2^2+\gamma\delta\sigma_1^2}{\alpha^2\sigma_2^2+\delta^2\sigma_1^2}\Big)\Big)^2+\Big(\frac{\alpha\gamma-\beta\delta}{\sqrt{\alpha^2\sigma_2^2+\delta^2\sigma_1^2}}\Big)^2\Big\}\Big\}$$

$$=\phi\Big(\sqrt{\frac{\alpha^2}{\sigma_1^2}+\frac{\delta^2}{\sigma_2^2}}\Big(x-\frac{\alpha\beta\sigma_2^2+\gamma\delta\sigma_1^2}{\alpha^2\sigma_2^2+\delta^2\sigma_1^2}\Big)\Big).\phi\Big(\frac{\alpha\gamma-\beta\delta}{\sqrt{\alpha^2\sigma_2^2+\delta^2\sigma_1^2}}\Big) \tag{2}$$

Now integrating both sides of (6) with respect to $x$ over the range $(-\infty,\infty)$, we have,

$$\int_{-\infty}^{\infty}\phi\Big(\frac{\alpha x-\beta}{\sigma_1}\Big).\phi\Big(\frac{\gamma-\delta x}{\sigma_2}\Big)dx$$

$$=\phi\Big(\frac{\alpha\gamma-\beta\delta}{\sqrt{\alpha^2\sigma_2^2+\delta^2\sigma_1^2}}\Big)\int_{-\infty}^{\infty}\phi\Big(\sqrt{\frac{\alpha^2}{\sigma_1^2}+\frac{\delta^2}{\sigma_2^2}}\Big(x-\frac{\alpha\beta\sigma_2^2+\gamma\delta\sigma_1^2}{\alpha^2\sigma_2^2+\delta^2\sigma_1^2}\Big)\Big)dx$$

$$=\frac{1}{\sqrt{\frac{\alpha^2}{\sigma_1^2}+\frac{\delta^2}{\sigma_2^2}}}\phi\Big(\frac{\alpha\gamma-\beta\delta}{\sqrt{\alpha^2\sigma_2^2+\delta^2\sigma_1^2}}\Big) \tag{3}$$

□

### Appendix B

**Result 1.** *Using the model (1-3),*

$$P(y_i|\boldsymbol{Z}_{i,o}) = \sigma\Big(\frac{y_i\beta(\beta_0\boldsymbol{\beta}_x'\boldsymbol{X}_i+\beta_gG_i+\beta_mM_i+\beta_e\mu_0)}{\sqrt{\beta^2+y_i^2\beta_e^2+\sigma_2^2}}\Big) \tag{4}$$

*where* $\mu_0 = \gamma_0 + \gamma_g G_i + \gamma_m M_i$*, for each* $i \in S_{-E}$*, where* $S_{-E}$ *be the set of* $n_2$ *individuals for whom gene expression data are not available.*

*Proof.* We write, $\boldsymbol{Z}_i = (\boldsymbol{Z}'_{i,o}, \boldsymbol{Z}'_{i,m})'$ for all $i \in S_{-E}$ where the suffixes '$o$' and '$m$' denote observed and missing parts. So, here $\boldsymbol{Z}_{i,o} = (1, \boldsymbol{X}'_i, G_i, M_i)'$ and $\boldsymbol{Z}_{i,m} = E_i$ for all $i \in S_{-E}$. Note that $\mathscr{C}(S_{-E}) = n_2$ where $\mathscr{C}(A)$ denotes the cardinality of a set $A$.

Now for each $i \in S_{-E}$, the observations on phenotype, genotype, covariates and methylation are known but gene expression values are unknown. Hence, we have for each $i \in S_{-E}$,

$$
\begin{aligned}
P(y_i|\boldsymbol{Z}_{i,o}) &= \int_{Z_{i,m}} P(y_i|\boldsymbol{Z}_{i,o}, \boldsymbol{Z}_{i,m}) P(\boldsymbol{Z}_{i,m}|\boldsymbol{Z}_{i,o}) dZ_{i,m} \\
&= \int_{E_i} \sigma(y_i(\beta_0 + \boldsymbol{\beta}'_x \boldsymbol{X}_i + \beta_g G_i + \beta_m M_i + \beta_e E_i)) \frac{1}{\sigma_2} \phi(\frac{E_i - \gamma_0 - \gamma_g G_i - \gamma_m M_i}{\sigma_2}) dE_i \\
&\approx \int_{E_i} \int_{u_i=-\infty}^{y_i(\boldsymbol{w}'\boldsymbol{Z}_i)} \phi(\frac{u}{\beta}) \frac{1}{\sigma_2} \phi(\frac{E_i - \mu_0}{\sigma_2}) du\, dE_i \quad \text{where, } \beta = \frac{\pi}{\sqrt{3}} \text{ [1]} \\
&= \int_{E_i} \int_{u_i=-\infty}^{y_i(\boldsymbol{w}'_o\boldsymbol{Z}_{i,o})} \phi(\frac{\nu + y_i\beta_e E_i}{\beta}) \frac{1}{\sigma_2} \phi(\frac{(E_i - \mu_0) y_i \beta_e}{\sigma_2 y_i \beta_e}) d\nu\, dE_i \quad \text{where, } u = \nu + \beta_e E_i y_i
\end{aligned}
$$

Now putting $x = E_i$, $\alpha = y_i\beta_e$, $\beta = \mu_0 y_i \beta_e$, $\gamma = \nu$, $\delta = -y_i\beta_i$, $\sigma_1 = \beta$, $\sigma_2 = \sigma_2 y_i \beta_e$ in Lemma 1, we have,

$$
\begin{aligned}
P(y_i|\boldsymbol{Z}_{i,o}) &= \frac{1}{\beta_e \sigma_2} \frac{1}{\sqrt{2\pi}\sqrt{\frac{y_i^2}{\beta^2} + \frac{1}{\beta_e^2\sigma_2^2}}} \int_{\nu=-\infty}^{y_i\boldsymbol{w}'_o\boldsymbol{Z}_{i,o}} e^{-\frac{1}{2}\frac{(\nu + \beta_e y_i \mu_0)^2}{\beta^2 + y_i^2\beta_e^2\sigma_2^2}} d\nu \\
&= \frac{1}{\sqrt{\frac{\beta^2 + y_i^2\beta_e^2\sigma_2^2}{\beta^2}}} \int_{\nu=-\infty}^{y_i\boldsymbol{w}'_o\boldsymbol{Z}_{i,o}} \phi(\frac{\nu + \beta_e y_i \mu_0}{\sqrt{\beta^2 + y_i^2\beta_e^2\sigma_2^2}}) d\nu \\
&= \int_{\nu'=-\infty}^{\frac{y_i\boldsymbol{w}'_o\boldsymbol{Z}_{i,o} + y_i\beta_e\mu_0}{\lambda}} \phi(\frac{\nu'}{\beta}) d\nu' \text{ where, } \nu' = \frac{\nu + \beta_e y_i \mu_0}{\lambda}\ ,\ \lambda = \sqrt{1 + \frac{y_i^2\beta_e^2\sigma_2^2}{\beta^2}} \\
&\approx \sigma(\frac{y_i\boldsymbol{w}'_o\boldsymbol{Z}_{i,o} + y_i\beta_e\mu_0}{\lambda}) \qquad\qquad [\text{using (4.8)}] \\
&= \sigma(\frac{y_i\beta(\beta_0 + \boldsymbol{\beta}'_x\boldsymbol{X}_i + \beta_g G_i + \beta_m M_i + \beta_e\mu_0)}{\sqrt{\beta^2 + y_i^2\beta_e^2\sigma_2^2}})
\end{aligned}
$$

□

**Result 2.** *Using the model (1-3), for each* $i \in S_{-M}$*,*

$$
P(y_i|\boldsymbol{Z}_i) = \sigma\Bigg(\frac{y_i\beta(\beta_0 + \boldsymbol{\beta}'_x\boldsymbol{X}_i + \beta_g G_i + \beta_e E_i + \beta_m \frac{(\alpha_0 + \alpha_g G_i)\sigma_2^2 + \gamma_m(E_i - \gamma_0 - \gamma_g G_i)\sigma_1^2}{\sigma_2^2 + \gamma_m^2\sigma_1^2})}{\sqrt{\beta^2 + \beta_m^2 \frac{\sigma_1^2\sigma_2^2}{\sigma_2^2 + \gamma_m^2\sigma_1^2}}}\Bigg) \tag{5}
$$

*where* $S_{-M}$ *is the set of* $n_3$ *individuals for whom no methylation data are available.*

*Proof.* From model (1-3), we have, $E(M_i) = \alpha_0 + \alpha_g G_i$, $V(M_i) = \sigma_1^2$. Now,

$$
\begin{aligned}
&E(E_i) = E_{M_i}E_{E_i|M_i}(E_i|M_i) = E_{M_i}(\gamma_0 + \gamma_g G_i + \gamma_m M_i) = \gamma_0 + \gamma_g G_i + \gamma_m(\alpha_0 + \alpha_g G_i) \\
&V(E_i) = E_{M_i}V_{E_i|M_i}(E_i|M_i) + V_{M_i}E_{E_i|M_i}(E_i|M_i) = \sigma_2^2 + V_{M_i}(\gamma_0 + \gamma_g G_i + \gamma_m M_i) = \sigma_2^2 + \gamma_m^2\sigma_1^2 \\
&Cov(M_i, E_i) = Cov(M_i, \gamma_0 + \gamma_g G_i + \gamma_m M_i + \epsilon_{2i}) = \gamma_m V(M_i) = \gamma_m\sigma_1^2 \\
&\rho_{M_i,E_i} = \frac{Cov(M_i, E_i)}{\sqrt{V(E_i)V(M_i)}} = \frac{\gamma_m\sigma_1^2}{\sqrt{\sigma_1^2(\sigma_2^2 + \gamma_m^2\sigma_1^2)}} = \frac{\gamma_m\sigma_1}{\sqrt{\sigma_2^2 + \gamma_m^2\sigma_1^2}}
\end{aligned}
$$

Hence, $P(E_i|M_i) = \frac{1}{\sigma_2}\phi(\frac{E_i-\gamma_0-\gamma_g G_i-\gamma_m M_i-\boldsymbol{\gamma}_x'\boldsymbol{X}_i}{\sigma_2})$.
Now, for each $i \in S_{-M}$,

$$\begin{aligned} P(y_i|\boldsymbol{Z}_{i,o}) &= \int_{Z_{i,m}} P(y_i|\boldsymbol{Z}_{i,o},\boldsymbol{Z}_{i,m})P(\boldsymbol{Z}_{i,m}|\boldsymbol{Z}_{i,o})dZ_{i,m} \\ &= \int_{M_i=-\infty}^{\infty} \sigma(y_i(\boldsymbol{w}'\boldsymbol{Z}_i))P(M_i|E_i)dM_i = \int_{M_i=-\infty}^{\infty} \sigma(y_i(\boldsymbol{w}'\boldsymbol{Z}_i))\frac{P(E_i|M_i)P(M_i)}{P(E_i)}dM_i \end{aligned} \tag{6}$$

Now, Denominator in (12) is

$$\begin{aligned} P(E_i) &= \int_{M_i} P(E_i,M_i)dM_i = \int_{M_i} P(E_i|M_i)P(M_i)dM_i \\ &= \frac{1}{\sigma_1\sigma_2}\int_{M_i}\phi\Big(\frac{E_i-\gamma_0-\gamma_g G_i-\gamma_m M_i}{\sigma_2}\Big)\phi\Big(\frac{M_i-\alpha_0-\alpha_g G_i}{\sigma_1}\Big)dM_i \end{aligned}$$

Now in Lemma 1, put $\alpha = 1$, $\beta = \alpha_0 + \alpha_g G_i$, $\sigma_1 = \sigma_1$, $\gamma = E_i - \gamma_0 - \gamma_g G_i$, $\delta = \gamma_m$, $\sigma_2 = \sigma_2$, and noting that $\phi(-x) = \phi(x)$, we have,

$$\begin{aligned} P(E_i) &= \frac{1}{\sigma_1\sigma_2}\frac{1}{\sqrt{\frac{1}{\sigma_1^2}+\frac{\gamma_m^2}{\sigma_2^2}}}.\phi\Big(\frac{E_i-\gamma_0-\gamma_g G_i-\gamma_m(\alpha_0+\alpha_g G_i)}{\sqrt{\sigma_2^2+\gamma_m^2\sigma_1^2}}\Big) \\ &= \frac{1}{\sqrt{\sigma_2^2+\gamma_m^2\sigma_1^2}}.\phi\Big(\frac{a\gamma_m-b}{\sqrt{\sigma_2^2+\gamma_m^2\sigma_1^2}}\Big) \end{aligned} \tag{7}$$

where $a = \alpha_0 + \alpha_g G_i$ and $b = E_i - \gamma_0 - \gamma_g G_i$

Now putting $p = \frac{a\sigma_2^2+\gamma_m b\sigma_1^2}{\sigma_2^2+\gamma_m^2\sigma_1^2}$, $q = \frac{1}{\sqrt{\frac{1}{\sigma_1^2}+\frac{\gamma_m^2}{\sigma_2^2}}} = \frac{\sigma_1\sigma_2}{\sigma_2^2+\gamma_m^2\sigma_1^2}$ and using Lemma 1 and (8), we can simplify the numerator of (12) as:

$$\begin{aligned} &\int_{M_i=-\infty}^{\infty}\sigma(y_i(\boldsymbol{w}'\boldsymbol{Z}_i))P(M_i|E_i)P(M_i)dM_i \\ &= \frac{1}{\sigma_1\sigma_2}\int_{M_i}\sigma(y_i\boldsymbol{w}'\boldsymbol{Z}_i)\phi\Big(\frac{E_i-\gamma_0-\gamma_g G_i-\gamma_m M_i}{\sigma_2}\Big)\phi\Big(\frac{M_i-\alpha_0-\alpha_g G_i}{\sigma_1}\Big)dM_i \\ &= \frac{1}{\sigma_1\sigma_2}\int_{M_i}\sigma(y_i\boldsymbol{w}'\boldsymbol{Z}_i)\phi\Big(\frac{a\gamma_m-b}{\sqrt{\sigma_2^2+\gamma_m^2\sigma_1^2}}\Big)\phi\Big(\sqrt{\frac{1}{\sigma_1^2}+\frac{\gamma_m^2}{\sigma_2^2}}(M_i-\frac{a\sigma_2^2+\gamma_m^2 b\sigma_1^2}{\sigma_2^2+\gamma_m^2\sigma_1^2})\Big)dM_i \\ &\approx \frac{1}{\sigma_1\sigma_2}\phi(\frac{a\gamma_m-b}{\sqrt{\sigma_2^2+\gamma_m^2\sigma_1^2}})\int_{M_i}\int_{u=-\infty}^{y_i\boldsymbol{w}'\boldsymbol{Z}_i}\phi(\frac{u}{\beta})\phi(\frac{M_i-p}{q})\,du\,dM_i \\ &= \frac{1}{\sigma_1\sigma_2}\frac{1}{\beta_m}\phi(\frac{a\gamma_m-b}{\sqrt{\sigma_2^2+\gamma_m^2\sigma_1^2}})\int_{M_i^*}\int_{\nu=-\infty}^{y_i(\beta_0+\boldsymbol{\beta}_x'\boldsymbol{X}_i+\beta_g G_i+\beta_e E_i)}\phi(\frac{\nu+y_iM_i^*}{\beta})\phi(\frac{M_i^*-\beta_m p}{\beta_m q})\,d\nu\,dM_i^* \\ &\text{where } u = \nu + y_i\beta_m M_i,\ M_i^* = \beta_m M_i \\ &= \frac{\beta q}{\sigma_1\sigma_2\sqrt{\beta^2+\beta_m^2q^2}}\phi(\frac{a\gamma_m-b}{\sqrt{\sigma_2^2+\gamma_m^2\sigma_1^2}})\int_{\nu=-\infty}^{y_i(\beta_0+\boldsymbol{\beta}_x'\boldsymbol{X}_i+\beta_g G_i+\beta_e E_i)}\phi(\frac{\nu+y_i\beta_m p}{\sqrt{\beta^2+\beta_m^2q^2}})d\nu \\ &= \frac{q}{\sigma_1\sigma_2}\phi(\frac{a\gamma_m-b}{\sqrt{\sigma_2^2+\gamma_m^2\sigma_1^2}})\int_{\nu'=-\infty}^{\frac{y_i\beta(\beta_0+\boldsymbol{\beta}_x\boldsymbol{X}_i+\beta_g G_i+\beta_e E_i+\beta_m p)}{\sqrt{\beta^2+\beta_m^2q^2}}}\phi(\frac{\nu'}{\beta})d\nu',\ \text{where } \frac{\nu'}{\beta} = \frac{\nu+y_i\beta_m p}{\sqrt{\beta^2+\beta_m^2q^2}} \\ &\approx \frac{1}{\sqrt{\sigma_2^2+\gamma_m^2\sigma_1^2}}\phi(\frac{a\gamma_m-b}{\sqrt{\sigma_2^2+\gamma_m^2\sigma_1^2}})\sigma(\frac{y_i\beta(\beta_0+\boldsymbol{\beta}_x'\boldsymbol{X}_i+\beta_g G_i+\beta_e E_i+\beta_m p)}{\sqrt{\beta^2+\beta_m^2q^2}})\ \text{[using (8)]} \end{aligned} \tag{8}$$

Therefore, using (13) and (14) in (12), we have,

$$\begin{aligned}P(y_i|\boldsymbol{Z}_{i,o}) &= \int_{M_i=-\infty}^{\infty} \sigma(y_i(\boldsymbol{w}'\boldsymbol{Z}_i))\frac{P(E_i|M_i)P(M_i)}{P(E_i)}dM_i \\ &\approx \sigma\Big(\frac{y_i\beta(\beta_0+\boldsymbol{\beta}_x'\boldsymbol{X}_i+\beta_g G_i+\beta_e E_i+\beta_m p)}{\sqrt{\beta^2+\beta_m^2 q^2}}\Big) \\ &= \sigma\Big(\frac{y_i\beta(\beta_0+\boldsymbol{\beta}_x'\boldsymbol{X}_i+\beta_g G_i+\beta_e E_i+\beta_m \frac{(\alpha_0+\alpha_g G_i)\sigma_2^2+\gamma_m(E_i-\gamma_0-\gamma_g G_i)\sigma_1^2}{\sigma_2^2+\gamma_m^2\sigma_1^2})}{\sqrt{\beta^2+\beta_m^2\frac{\sigma_1^2\sigma_2^2}{\sigma_2^2+\gamma_m^2\sigma_1^2}}}\Big) \end{aligned} \tag{9}$$

□

**Result 3.** *Under the model (1-3), for each $i \in S_{-(E,M)}$,*

$$P(y_i|\boldsymbol{Z}_{i,o}) = \sigma(\frac{y_i\beta(p_1(\beta_0+\boldsymbol{\beta}_x X_i+\beta_g G_i)+\beta_m a\sigma_2^2-\beta_m\gamma_m\sigma_1^2(\gamma_0+\gamma_g G_i)+\delta_3)}{\sqrt{\delta_2^2+\delta_4^2}}) \tag{10}$$

*where $S_{-(E,M)}$ is the set of $n_4$ individuals for whom both expression and methylation data are missing.*

*Proof.* For each $i \in S_{-(E,M)}$,

$$\begin{aligned}P(y_i|\boldsymbol{Z}_{i,o}) &= \int\limits_{Z_{i,m}} P(y_i|\boldsymbol{Z}_{i,o},\boldsymbol{Z}_{i,m})P(\boldsymbol{Z}_{i,m}|\boldsymbol{Z}_{i,o})\,dZ_{i,m} \\ &= \int\limits_{M_i=-\infty}^{\infty}\int\limits_{E_i=-\infty}^{\infty} \sigma(y_i(\boldsymbol{w}'\boldsymbol{Z}_i))P(E_i|M_i)P(M_i)\,dE_i\,dM_i \\ &= \int\limits_{M_i=-\infty}^{\infty}\int\limits_{E_i=-\infty}^{\infty} \Big\{\sigma(y_i(\boldsymbol{w}'\boldsymbol{Z}_i))\frac{1}{\sigma_2}\phi(\frac{E_i-\gamma_0-\gamma_g G_i-\gamma_m M_i-\boldsymbol{\gamma}_x'\boldsymbol{X}_i}{\sigma_2}) \\ &\qquad \times\frac{1}{\sigma_1}\phi(\frac{M_i-\alpha_0-\alpha_g G_i-\boldsymbol{\alpha}_x'\boldsymbol{X}_i}{\sigma_1})\Big\}dE_i dM_i \\ &\approx \int\limits_{M_i=-\infty}^{\infty}\int\limits_{E_i=-\infty}^{\infty}\int\limits_{u=-\infty}^{y_i\boldsymbol{w}'\boldsymbol{Z}_i} \phi(\frac{u}{\beta})\frac{1}{\sigma_2}\phi(\frac{E_i-\gamma_0-\gamma_g G_i-\gamma_m M_i}{\sigma_2})\frac{1}{\sigma_1}\phi(\frac{M_i-\alpha_0-\alpha_g G_i}{\sigma_1})du dE_i dM_i \end{aligned}$$

where, $\beta = \frac{\pi}{\sqrt{3}}$ [1]

$$\begin{aligned}&= \int\limits_{M_i}\int\limits_{E_i}\int\limits_{\nu=-\infty}^{y_i(\beta_0+\boldsymbol{\beta}_x' X_i+\beta_g G_i)} \Big\{\frac{1}{\sigma_1\sigma_2}\phi(\frac{\nu+y_i\beta_e E_i+y_i\beta_m M_i}{\beta})\phi(\frac{E_i-\gamma_0-\gamma_g G_i-\gamma_m M_i}{\sigma_2}) \\ &\qquad \times\phi(\frac{M_i-\alpha_0-\alpha_g G_i}{\sigma_1})\Big\}\,d\nu\,dE_i\,dM_i, \text{ where, } u=\nu+y_i\beta_e E_i+y_i\beta_m M_i \end{aligned} \tag{11}$$

Now in Lemma 1, put $x = E_i$, $\alpha = 1$, $\beta = \gamma_0+\gamma_g G_i+\gamma_m M_i$, $\sigma_1 = \sigma_2$, $\gamma = \nu + y_i\beta_m M_i$, $\delta = -y_i\beta_e$, and $\sigma_2 = \beta = \frac{\pi}{\sqrt{3}}$, to get,

$$\begin{aligned}&\int\limits_{E_i=-\infty}^{\infty} \phi(\frac{\nu+y_i\beta_e E_i+y_i\beta_m M_i}{\beta})\phi(\frac{E_i-\gamma_0-\gamma_g G_i-\gamma_m M_i}{\sigma_2})\,dE_i \\ &= \frac{1}{\sqrt{\frac{1}{\sigma_2^2}+\frac{y_i^2\beta_e^2}{\beta^2}}}\phi(\frac{\nu+y_i\beta_m M_i+y_i\beta_e(\gamma_0+\gamma_g G_i+\gamma_m M_i)}{\sqrt{\beta^2+y_i^2\beta_e^2\sigma_2^2}}) \end{aligned} \tag{12}$$

Again putting $x = M_i$, $\alpha = 1$, $\beta = \alpha_0 + \alpha_g G_i$, $\sigma_1 = \sigma_1$, $\gamma = \nu + y_i\beta_e(\gamma_0 + \gamma_g G_i)$, $\delta = -(y_i\beta_e\gamma_m + y_i\beta_m$, $\sigma_2 = \sqrt{\beta^2 + y_i^2\beta_e^2\sigma_2^2}$, we have,

$$\begin{aligned}
&\frac{1}{\sqrt{\frac{1}{\sigma_2^2}+\frac{y_i^2\beta_e^2}{\beta^2}}}\int\limits_{M_i=-\infty}^{\infty}\Bigg\{\phi(\frac{M_i-\alpha_0-\alpha_h G_i)}{\sigma_1}\\
&\qquad\times\phi(\frac{\nu+y_i\beta_m M_i+y_i\beta_e(\gamma_0+\gamma_g G_i+\gamma_m M_i)}{\sqrt{\beta^2+y_i^2\beta_e^2\sigma_2^2}})\Bigg\}dM_i\\
&=\frac{1}{\sqrt{\frac{1}{\sigma_2^2}+\frac{y_i^2\beta_e^2}{\beta^2}}}\frac{1}{\sqrt{\frac{1}{\sigma_1^2}+\frac{y_i^2(\beta_e\gamma_m+\beta_m)^2}{\beta^2+y_i^2\beta_e^2\sigma_2^2}}}\\
&\quad\times\phi(\frac{\nu+y_i\beta_e(\gamma_0+\gamma_g G_i)+(\alpha_0+\alpha_g G_i)(y_i\beta_e\gamma_m+y_i\beta_m)}{\sqrt{\beta^2+y_i^2\beta_e^2\sigma_2^2+y_i^2(\beta_e\gamma_m+\beta_m)^2\sigma_1^2}})\\
&=\frac{1}{\sqrt{\frac{1}{\sigma_2^2}+\frac{y_i^2\beta_e^2}{\beta^2}}}\frac{1}{\sqrt{\frac{1}{\sigma_1^2}+\frac{y_i^2(\beta_e\gamma_m+\beta_m)^2}{\beta^2+y_i^2\beta_e^2\sigma_2^2}}}\phi(\frac{\nu+y_i\beta_e d+a(y_i\beta_e\gamma_m+y_i\beta_m)}{\sqrt{\beta^2+y_i^2\beta_e^2\sigma_2^2+y_i^2(\beta_e\gamma_m+\beta_m)^2\sigma_1^2}})\\
&=\frac{\beta\sigma_1\sigma_2}{\delta_1}\phi(\frac{\nu+y_i\beta_e d+a(y_i\beta_e\gamma_m+y_i\beta_m)}{\delta_1})
\end{aligned}\tag{13}$$

where $a = (\alpha_0 + \alpha_g G_i)$, $d = (\gamma_0 + \gamma_g G_i)$ and $\delta_1^2 = \beta^2 + y_i^2\beta_e^2\sigma_2^2 + y_i^2(\beta_e\gamma_m + \beta_m)^2\sigma_1^2$.

$$\begin{aligned}
P(y_i|\boldsymbol{Z}_{i,o}) &= \int\limits_{\nu=-\infty}^{y_i\boldsymbol{w}_o'\boldsymbol{Z}_{i,o}} \frac{1}{\sigma_1\sigma_2}\frac{\beta\sigma_1\sigma_2}{\delta_1}\phi(\frac{\nu+y_i\beta_e d+a(y_i\beta_e\gamma_m+y_i\beta_m)}{\delta_1})\,d\nu\\
&=\sigma(\frac{\beta y_i(\beta_0+\beta_g G_i+\boldsymbol{\beta}_x'\boldsymbol{X}_i+\beta_e(\gamma_0+\gamma_g G_i)+(\alpha_0+\alpha_g G_i)(\beta_e\gamma_m+\beta_m))}{\sqrt{\beta^2+y_i^2\beta_e^2\sigma_2^2+y_i^2(\beta_e\gamma_m+\beta_m)^2\sigma_1^2}})
\end{aligned}\tag{14}$$

□

Table S1: List of significant TiMEG genes associated with TSC disease

| | | | | | | | | | |
|---|---|---|---|---|---|---|---|---|---|
| ACACA | ACSS2 | ACTR3C | AKAP12 | ALDH1L1 | AMOTL2 | ANKRD36B | ANO7 | APP | ARHGEF16 |
| ARSG | ARTN | ASAP1 | ASPA | ASPRV1 | ATG9B | ATP1B2 | ATP6V1C2 | BCOR | BEND6 |
| C1GALT1 | C3orf35 | C3orf67 | C4orf19 | C4orf3 | C5orf22 | C8orf31 | CASQ1 | CBFA2T2 | CCDC149 |
| CCDC7 | CEBPG | CHST10 | CHST13 | CLCC1 | CLDN15 | CNGB3 | CNTFR | CNTNAP1 | CPD |
| CRB2 | CREB5 | CRIM1 | CSNK1G2 | CUL9 | DCBLD1 | DCP2 | DLX1 | DLX3 | DNAH17 |
| DPY19L1 | DPYSL5 | DRD4 | ECT2L | EFCAB1 | EFNA2 | ELOVL7 | EMX2OS | EPDR1 | ERCC6 |
| EXOC2 | FAAH2 | FAM110A | FGD5 | FGFR2 | FHDC1 | FOXN2 | FSCN2 | GNG4 | GPM6A |
| HIF3A | HIST1H4C | HTR2C | HTT | IL1B | IQCE | JAK3 | KANK3 | KCND3 | KCNQ1OT1 |
| KIAA1109 | KLHDC8A | KLHL18 | LAMC2 | LANCL2 | LDLRAP1 | LIFR | LLGL2 | LMNB1 | LOC100270746 |
| LOC407835 | LOC441455 | LOC730101 | LOC90246 | LRP1B | MAFF | MAGI3 | MAP4K4 | MMRN2 | MORN1 |
| MYH10 | MYH15 | MYLK2 | NAV1 | NCK2 | NEBL | NIPSNAP3B | NLRP14 | NMT1 | NOLC1 |
| NPR3 | NT5M | OBSCN | PCBP4 | PMEPA1 | PRKG1 | PRRT4 | PSAT1 | PTK6 | PTPDC1 |
| RAP2C | RBMS3 | RESP18 | RNF103 | RNF207 | RRP15 | SEL1L3 | SERPINF1 | SGSM2 | SIPA1L3 |
| SLC16A8 | SLC27A6 | SLC29A1 | SLC7A11 | SMAD7 | SMPDL3B | SMTNL2 | SNTG2 | SNX31 | SPRY1 |
| SPTBN1 | SRI | SRPK3 | STK25 | STX11 | SULF2 | TACO1 | TBC1D12 | TBC1D14 | TBX1 |
| TCEA2 | TG | THSD7A | TOX2 | TRAPPC2 | TRAPPC9 | TSC1 | TSC22D4 | USH2A | VILL |
| VIPR2 | VPS13D | WDR27 | WIPF1 | WNT2B | WWC2 | ZDHHC11 | ZNF239 | ZNF275 | ZNF876P |

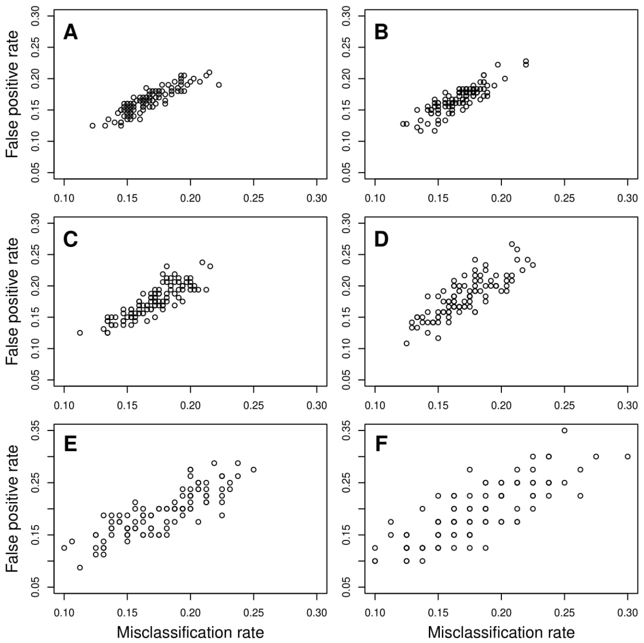


Figure $S1$: Plot of Misclassification rate vs False positive rate (1-Specificity) for only methylation missing. Figure S1A depicts no missing data scenario while Figures S1B-S1F respectively depict 10%, 20%, 40%, 60% and 80% only methylation data missing scenarios

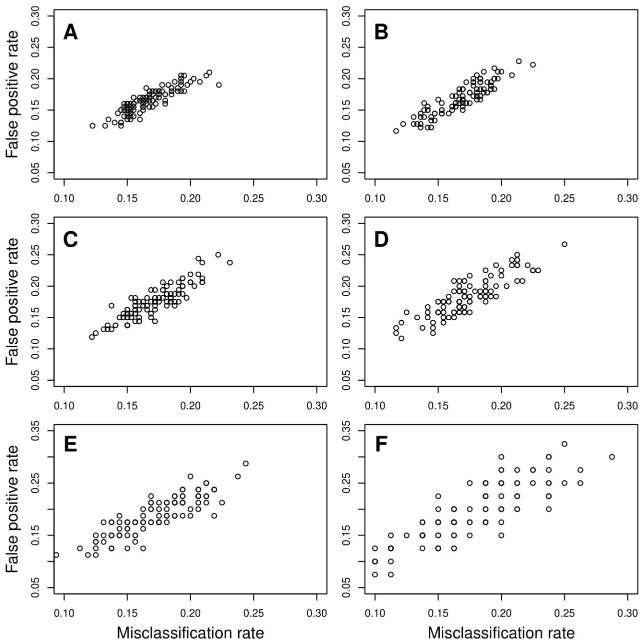


Figure $S2$: Plot of Misclassification rate vs False positive rate (1-Specificity) for both missing. Figure S2A depicts no missing data scenario while Figures S2B-S2F respectively depict 10%, 20%, 40%, 60% and 80% of both gene expression and methylation data missing scenarios

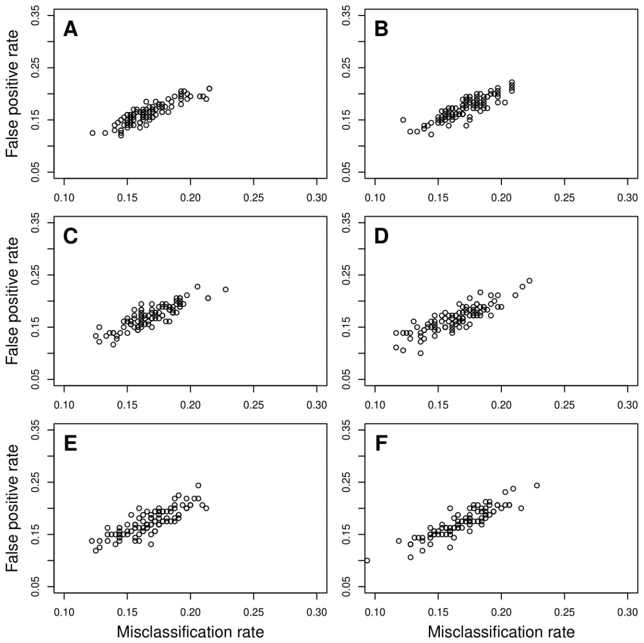


Figure $S3$: Plot of Misclassification rate vs False positive rate (1-Specificity) for miscellaneous missing. Figure S3A depicts no missing data scenario; Figure S3B depicts 5% individuals with methylation missing and another 5% with both gene expression and methylation missing but none of the individuals have only gene expression missing; similarly in Figure S3C the percentages of missing only methylation, both omics and only gene expression are respectively 0%, 5%, 5%; in Figures S3D-S3F these percentages are (5,0,5), (10,10,0), (5,10,5) respectively.
